# Human iPSC-derived hypertrophic chondrocytes reveal a mutation-specific unfolded protein response in chondrodysplasias

**DOI:** 10.1101/2020.05.19.103960

**Authors:** Yann Pretemer, Shunsuke Kawai, Makoto Watanabe, Sanae Nagata, Megumi Nishio, Sakura Tamaki, Cantas Alev, Jing-Yi Xue, Zheng Wang, Kenichi Fukiage, Masako Tsukanaka, Tohru Futami, Shiro Ikegawa, Junya Toguchida

## Abstract

Chondrodysplasias are hereditary diseases caused by mutations in the components of growth cartilage. Although the unfolded protein response (UPR) has been identified as a key disease mechanism in mouse models, no suitable *in vitro* system has been reported to analyze the pathology in humans. Here, utilizing human chondrodysplasia-specific iPSCs, we examined the UPR caused by mutations in *MATN3* or *COL10A1*. In growth plate-like structures formed from iPSC-derived sclerotome *in vivo*, the hypertrophic zone was disrupted, and induced hypertrophic chondrocytes *in vitro* showed varying levels of ER stress depending on the mutation. Autophagy inducers and chemical chaperones succeeded in reducing ER stress only in some mutants, while transcriptome analysis revealed many mutation-specific changes in genes involved in apoptosis, metabolism, and protein trafficking. In this way, our system has allowed the precise evaluation of the UPR caused by each mutation, opening up new avenues for treatment of individual chondrodysplasia patients.

## Introduction

Chondrodysplasias are hereditary cartilage disorders, which often manifest by early childhood as mild to severe skeletal abnormalities due to mutations in the components of growth cartilage. In multiple epiphyseal dysplasia (MED; OMIM #607078) and metaphyseal chondrodysplasia type Schmid (MCDS; OMIM #156500), short-limbed dwarfism and deformities of the hips or knees are commonly observed (Czarny-Ratajczak et al., 2001; Mäkitie et al., 2005), but large variations in the skeletal phenotype, disease severity, and types of mutations between patients have made these diseases difficult to research and treat. In order to overcome the obstacles posed by this heterogeneity and deepen our understanding of chondrodysplasias, it is imperative to obtain relevant patient samples and establish accurate disease models.

However, as it is ethically questionable to obtain samples from patients’ growth plates, most biopsies are taken from the iliac crest, which provides only limited information. Therefore, much of our understanding of chondrodysplasia disease mechanisms has come from studies of animal models. Mutations in the MATN3 vWFa and COL10A1 NC1 domains, which respectively cause MED and MCDS, have been reported to disrupt folding and oligomerization *in vitro* (Cotterill et al., 2005; Wilson et al., 2005), suggesting a gain-of-function effect that has been further supported by *MATN3* or *COL10A1* knockout mice having no significant skeletal dysplasia (Kwan et al., 1997; van der Weyden et al., 2006). In contrast, model mice with the p.V194D mutation in *Matn3*, known to cause MED in humans, showed short-limbed dwarfism and structural disruption of the growth plate with decreased chondrocyte proliferation and increased apoptosis (Leighton et al., 2007). Similarly, model mice with the MCDS-causing p.N617K mutation in *Col10a1* also showed short-limbed dwarfism, but the growth plate had an extended hypertrophic zone without decrease in chondrocyte proliferation (Rajpar et al., 2009). In both models, ER stress was detected in growth plate chondrocytes as a result of intracellular accumulation of MATN3 or COL10A1, indicating that the unfolded protein response (UPR) is a key event in these diseases. However, these results were obtained using homozygous mice, with heterozygotes having no or almost no phenotype. This is despite most MED- and MCDS-causing mutations being heterozygous with autosomal dominant inheritance in humans (Mortier et al., 2019). Due to these species differences, models that more closely reflect the pathology in humans are required.

Recently, disease-specific iPSCs have emerged as a powerful tool to further our understanding of human hereditary diseases and screen for candidate drugs. For example, the clinical phenotype of type II collagenopathies, a subgroup of chondrodysplasias, has been recapitulated *in vitro* using patient-derived iPSCs (Okada et al., 2015). Therefore, in this study, we aimed to apply this approach to MED and MCDS. As these two chondrodysplasias mainly or partly affect hypertrophic chondrocytes, a robust and efficient method for the induction of such late-stage chondrocytes from iPSCs is required. Since such a method has not yet been established, we first developed a 3D culture protocol to derive hypertrophic chondrocytes, which we characterized by cellular morphology and gene expression. Then, using both patient-derived and artificially mutated iPSC lines with heterozygous *MATN3* or *COL10A1* mutations, we applied our protocol to the *in vitro* recapitulation of MED and MCDS phenotypes. This enabled us to demonstrate a phenotype in humans that is similar to previous observations in homozygous model mice, including ER stress caused by the intracellular accumulation of the affected protein, the effects of which we further confirmed in an *in vivo* model of the growth plate. Comparison with isogenic controls enabled us to assess the contribution of each mutation to the phenotype and determine mutation-specific differences at the transcriptional level and in response to drugs, opening up new avenues to explore for the treatment of chondrodysplasias.

## Results

### Differentiation of iPSCs into hypertrophic chondrocytes through the sclerotome

In order to model chondrodysplasias *in vitro*, we first developed a protocol of differentiating hypertrophic chondrocytes from the wild type 414C2 iPSC line (Okita et al., 2011) in serum-free conditions. We combined previously reported protocols of sclerotome induction (Matsuda et al., 2020) and chondrogenic induction (Umeda et al., 2012), with slight modifications including the addition of the thyroid hormone T3, which has been reported to promote hypertrophic maturation (Mueller and Tuan, 2008). After sclerotome induction (SI), hypertrophic induction (HI) was performed in 3D culture for up to 70 days (Figure 1A).

**Figure 1.**
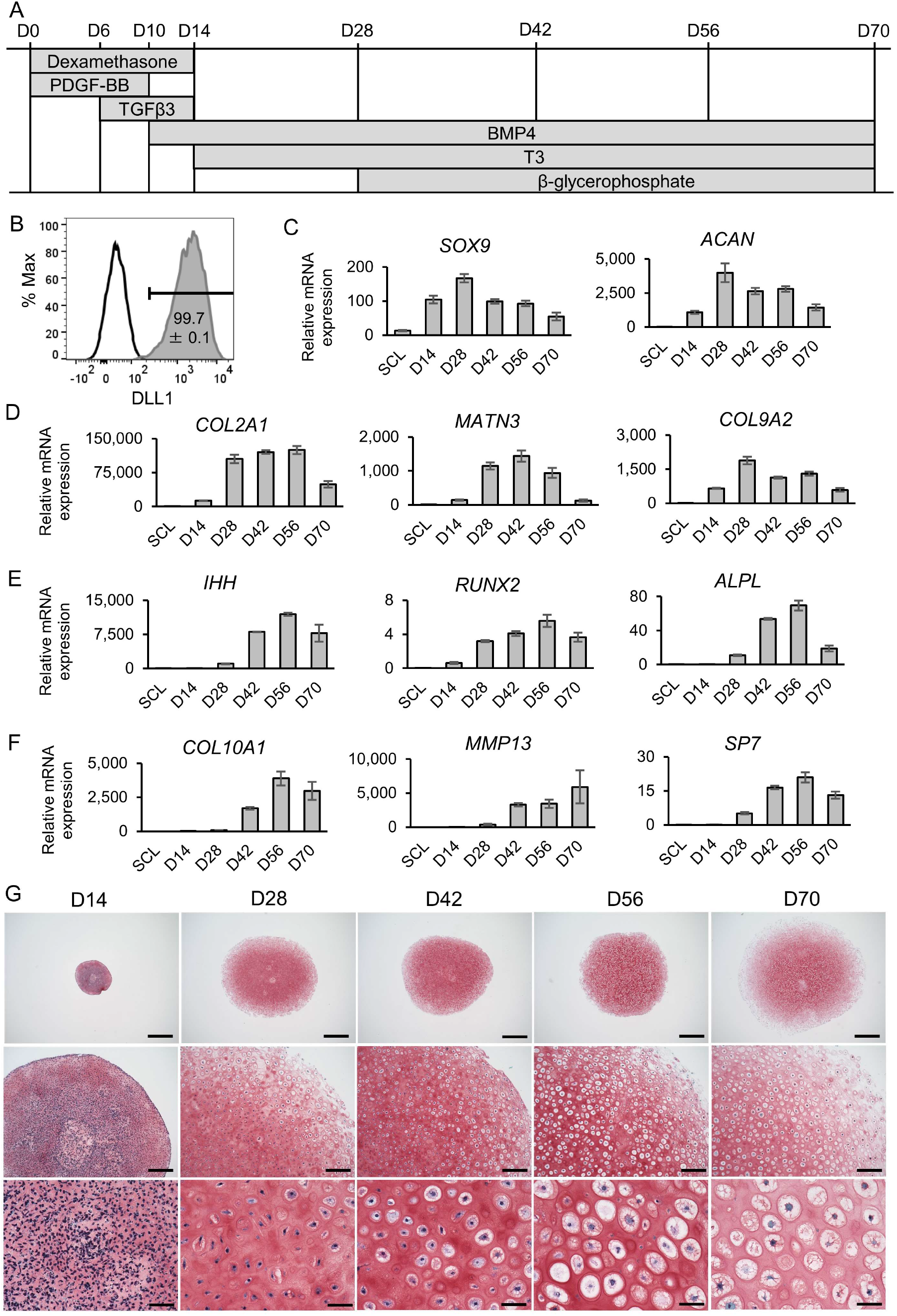
Differentiation of hypertrophic chondrocytes from iPSCs. (A) Protocol of hypertrophic induction (HI) in 3D pellet culture from sclerotome (SCL) cells. (B) Representative result of DLL1-positive cells (compared to isotype control) on day 2 of sclerotome induction (SI) with the mean and SEM (standard error of the mean) of four biological replicates displayed. (C-F) Expression of early (C), proliferating (D), pre-hypertrophic (E) and hypertrophic (F) chondrocyte markers over time from SCL on day −1 to day 70 of HI. Values are relative to the chondroblastic osteosarcoma cell line ANOS, which stably expresses these markers, and are shown as mean±SEM (n=4 from biologically independent experiments). (G) Representative result of Safranin O staining of pellets from day 14 to 70 of HI. Similar results were obtained in three biologically independent experiments. Scale bars, (top) 1 mm, (middle) 200 μm, (bottom) 50 μm. All results shown are from experiments using the 414C2 wild type iPSC line. See also Figure S1.

This protocol required no cell sorting, as FACS analysis at the presomitic mesoderm (PSM) stage during SI showed almost 100% of cells were positive for the PSM marker DLL1 (Figure 1B). The expression of early chondrocyte markers was detected from day 14 of HI, peaking at day 28 and declining again thereafter (Figure 1C). Proliferating and pre-hypertrophic chondrocyte markers increased from day 28 and declined by day 70 (Figures 1D and 1E), while hypertrophic markers mostly appeared on day 42 (Figure 1F). The pellet dramatically increased in size from 1 mm on day 14 to 4-5 mm on day 28, stabilizing thereafter (Figure 1G). On day 28, cells resembled proliferating chondrocytes in the pellet interior, with only a thin layer of cells with a pre-hypertrophic morphology in the periphery of the pellet, but by day 56, a hypertrophic morphology was detected throughout the pellet. The cartilage matrix stained with Safranin O from day 14 and was disintegrating in the periphery of the pellet by day 70. The expression of chondrocyte markers and the change in chondrocyte morphology during HI were similar in the 1231A3 iPSC line (Nakagawa et al., 2014) (Figures S1A-S1F).

### Patient analysis and establishment of *COL10A1* and *MATN3* mutant iPSC lines

iPSC lines were established from one MED patient with a previously reported heterozygous *MATN3* c.359C>T (p.T120M) mutation (Jackson et al., 2004), one MCDS patient with a novel heterozygous *COL10A1* c.1841_1841delT (p.L614Rfs*8) mutation, and one MCDS patient with a previously reported heterozygous *COL10A1* c.53G>A (p.G18E) mutation (Ikegawa et al., 1997). Karyotype analysis showed no chromosomal abnormalities in the two clones from each patient (Figures S2A, S3A, and S4A). Each clone showed normal morphology, the presence of pluripotency markers, and the ability to differentiate into all three germ layers (Figures S2B-S2D, S3B-S3D, and S4B-S4D).

Radiological findings in the MED patient included bowing of the femora with genu varum, as well as mild platyspondyly and scoliosis (Figure S2E). The patient’s height at age 9 was 2.2 SD (standard deviations) below normal. The MATN3 T120M mutation was corrected in the rescued clone (Figure S2F). In addition to the MATN3 T120M mutation, the SNP *MATN3* c.659T>C (p.V220A), which has been reported in both MED patients and normal controls (Kim et al., 2011), was also detected in the MED patient’s healthy allele (Figure S2G). To further analyze the pathology of *MATN3* mutations, a heterozygous *MATN3* c.626G>C (p.R209P) mutation was created in 414C2 iPSCs (Figure S2H). This mutation has been previously reported in MED, causing genu valgum but no dwarfism (Kim et al., 2011).

MCDS patient #1, who had the COL10A1 L614Rfs*8 mutation, showed typical a MCDS phenotype by age 2, with radiological findings including metaphyseal flaring and coxa vara (Figure S3E). The mutation was corrected in the rescued clone (Figure S3F). To test whether nonsense-mediated decay (NMD) occurred as a result of the early stop codon caused by the frameshift mutation, RNA of the patient-derived clone was reverse transcribed, amplified by PCR, and processed by restriction enzymes recognizing only the mutant or wild type allele (Figure S3G). Both alleles were present in the same amount, showing that the L614Rfs*8 mutation does not lead to NMD. MCDS patient #2, with the COL10A1 G18E mutation, has been previously described with radiological findings including widening of the physes, bowing of the femora, and coxa vara (Ikegawa et al., 1997). The mutation was corrected in the rescued clone (Figure S4E). Another heterozygous *COL10A1* mutant with the c.1798T>C (p.S600P) mutation was created using 414C2 iPSCs (Figure S4F). This mutation has been reported in an MCDS patient with short-limbed dwarfism, coxa vara, and metaphyseal abnormalities (Gregory et al., 2000).

### *COL10A1* and *MATN3* mutants differentiate into hypertrophic chondrocytes

We next assessed the ability of our mutant iPSC lines to differentiate into hypertrophic chondrocytes. At the PSM stage, mutants showed an equal or higher percentage of DLL1-positive cells (Figure 2A). On day 56 of HI, the expression of chondrocyte markers from various stages was similar in mutants compared to isogenic controls, with only a tendency of lower *IHH* expression observed (Figure S5A). *MATN3* expression in *MATN3* mutants was unchanged, but *COL10A1* expression tended to slightly decrease in *COL10A1* mutants (Figure 2B). Cartilage matrix production was not disrupted in mutants, as Safranin O staining and the pellet size were similar to isogenic controls (Figures 2C, S5B, and 2D). Despite the changes in *IHH* and *COL10A1* expression, the cell morphology was not different between mutants and controls, with both showing a hypertrophic morphology. The cell size showed no consistent changes for *COL10A1* or *MATN3* mutants (Figure 2E). Contrary to expectations, cell death was unchanged or actually decreased in mutants, indicating that *COL10A1* and *MATN3* mutations do not cause apoptosis in human cartilage (Figure 2F).

**Figure 2.**
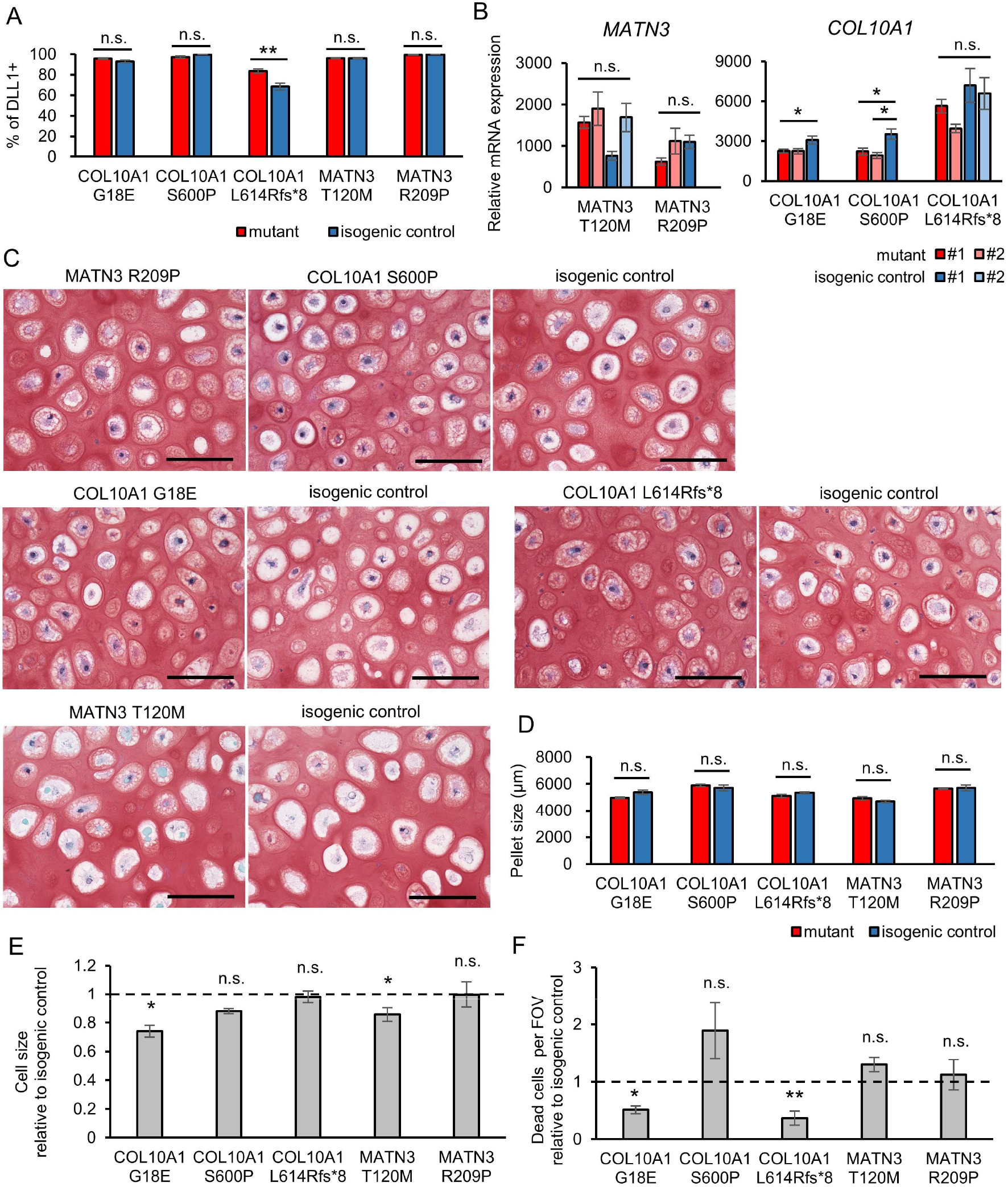
*COL10A1* and *MATN3* mutants show no disruption of chondrogenic differentiation. (A) Percentage of DLL1-positive cells on day 2 of SI. (B) mRNA expression of the heterozygously mutated gene, with values from a second clone also shown for each mutant and isogenic control (biologically independent sample number shown in Table S1). Values are relative to ANOS. Statistical analysis by ANOVA and post-hoc Tukey HSD. (C) Representative result of Safranin O staining of mutants (left) and their isogenic controls (right). Similar results were obtained in four biologically independent experiments. Scale bars, 100 μm. (D) Pellet size of each mutant and its isogenic control. Three technical replicates per biological replicate were measured. (E) Cell size of each mutant relative to the isogenic control, quantified from the inverse image of COL2A1 fluorescence. (F) TUNEL-positive cells per FOV (field of view) relative to the isogenic control of each mutant. All results except (A) are from day 56 of HI. Values are expressed as mean±SEM. Dotted lines indicate the value=1 of the isogenic controls in (E) and (F). Except where stated otherwise, the number of biological replicates is four and statistical analysis was performed using unpaired two-sided t-tests. (n.s. no significant difference, *p < 0.05, **p < 0.01, ***p < 0.001, ****p < 0.0001). See also Figures S2-S5.

### COL10A1 or MATN3 is retained intracellularly in mutants

Since MCDS and MED model mice harboring *COL10A1* or *MATN3* mutations accumulate the affected protein intracellularly, we examined whether this is also the case in humans. Indeed, immunostaining of the protein on day 56 of HI revealed the presence of intracellular aggregates in both *COL10A1* and *MATN3* mutants (Figures 3A, 3B, S5C, and S5D). These aggregates costained with the ER marker PDI, showing that they are retained within the ER. At the same time, extracellular COL10A1 or MATN3 was decreased in mutants. All mutants showed a significant increase of COL10A1 or MATN3 retention, which differed depending on the mutation and was particularly elevated in the MATN3 T120M mutant (Figures 3C and 3D).

**Figure 3.**
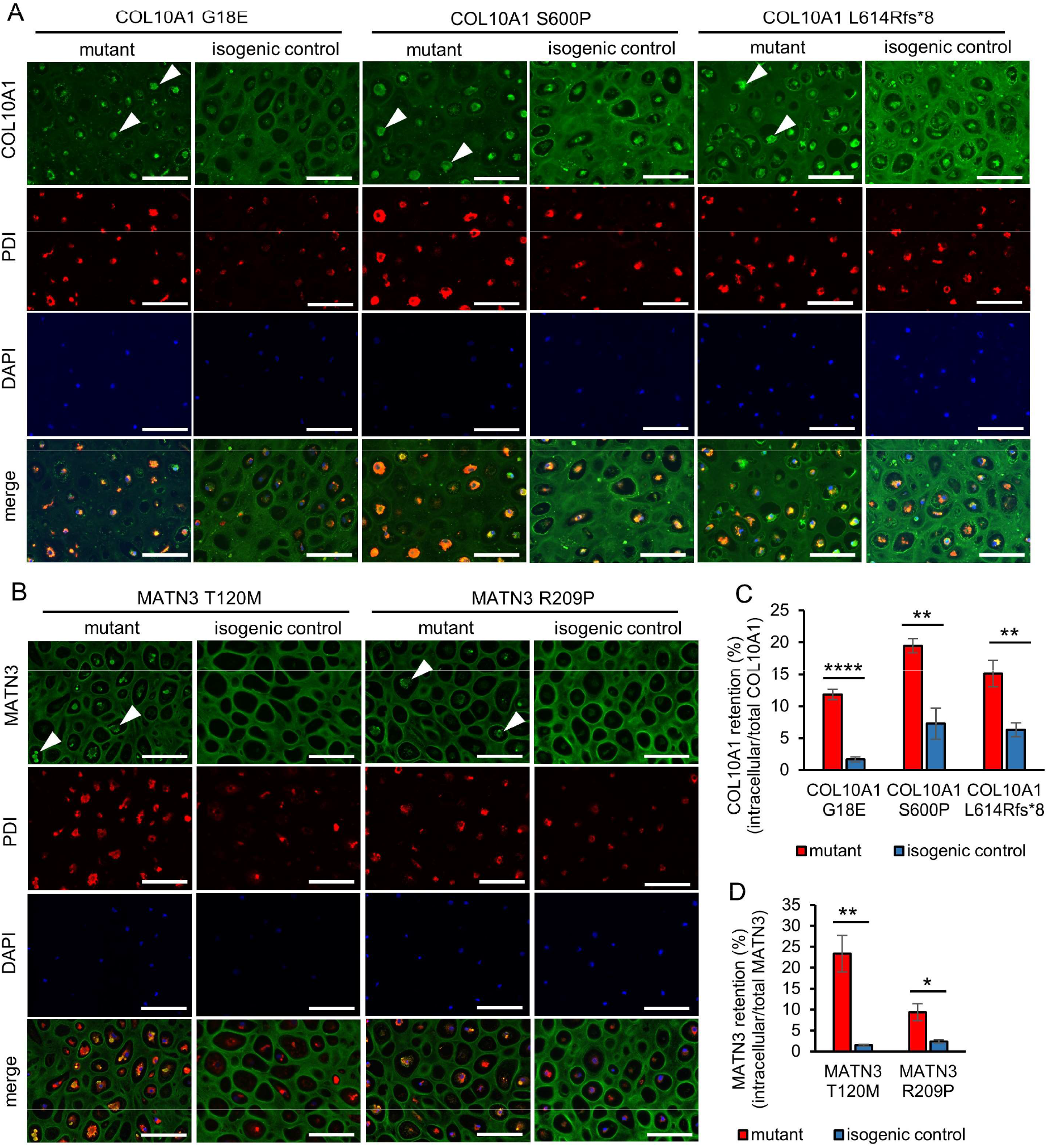
*COL10A1* and *MATN3* mutants retain COL10A1 or MATN3 within the ER. (A, B) Immunostaining of COL10A1 (A) or MATN3 (B) and PDI on day 56 of HI. Arrowheads indicate intracellular aggregates in mutants. Similar results were obtained in four biologically independent experiments. Scale bars, 100 μm. (C, D) Intracellular retention quantified from fluorescence intensity of COL10A1 (C) or MATN3 (D) co-staining with PDI (intracellular) divided by total fluorescence intensity in n=4 biological replicates on day 56 of HI. Values are expressed as mean±SEM. (n.s. no significant difference, *p < 0.05, **p < 0.01, ***p < 0.001, ****p < 0.0001 by unpaired twosided t-test). See also Figure S5.

### *COL10A1* and *MATN3* mutants show ER stress and enlargement of the ER

To test whether the intracellular accumulation of COL10A1 and MATN3 leads to ER stress in human cartilage, we looked at the expression of ER stress markers using qPCR. In all clones, one or more markers were significantly elevated at the mRNA level, with most clones also showing significant increases at the protein level (Figures 4A, S5E, and 4B). The MATN3 T120M mutant again showed the greatest increase of these markers, concordant with its high intracellular retention. Using transmission electron microscopy (TEM), we found a mildly to severely enlarged ER in all mutants (Figure 4C). Quantification of the PDI-positive areas confirmed these results, with all but the MATN3 R209P mutant showing a significant increase in ER size (Figure 4D). The COL10A1 S600P mutant, which showed the greatest increase, also had the most significantly increased ER stress markers of all *COL10A1* mutants. Surprisingly, the COL10A1 G18E mutant, which showed the most significant increase in the ER size, showed only slightly higher ER stress markers at the mRNA level and only a tendency at the protein level, indicating that the amount of ER stress does not necessarily correlate with ER size.

**Figure 4.**
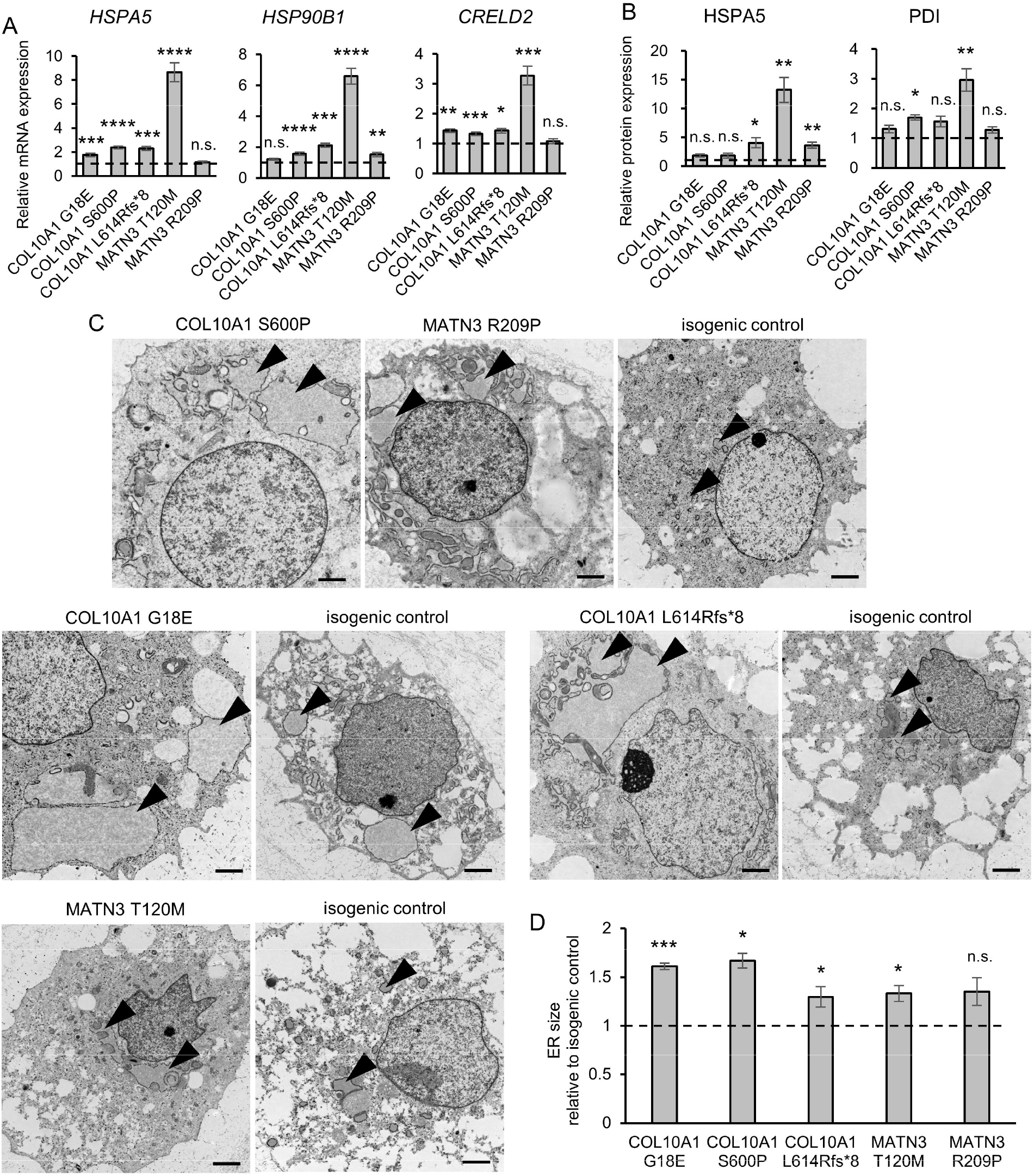
*COL10A1* and *MATN3* mutants show ER stress and a highly distended ER. (A) mRNA expression of ER stress markers (biologically independent sample number shown in Table S1). (B) Protein expression of ER stress markers by Simple Western from n=4 biologically independent experiments. (C) Transmission electron microscopy (TEM) of mutants (left) and their isogenic controls (right). Arrowheads indicate the ER. Scale bars, 2 μm. (D) ER size quantified from the area of PDI fluorescence from n=4 biologically independent experiments. All results are from day 56 of HI and are expressed as mean±SEM, relative to the respective isogenic control (dotted lines). Statistical analysis was performed by unpaired two-sided t-test (n.s. no significant difference, *p < 0.05, **p < 0.01, ***p < 0.001, ****p < 0.0001). See also Figure S5.

### Zonal disorganization and morphological abnormalities in growth plate-like structures

Since cell morphology and chondrocyte maturation markers in mutants showed little or no change in our *in vitro* system, we next assessed whether an *in vivo* environment may allow the detection of further differences. iPSC-derived sclerotome cells, which served as the chondroprogenitors in our *in vitro* model, were transplanted into immunodeficient mice to allow spontaneous development into tissues including cartilage and self-organizing growth plate-like structures (Matsuda et al., 2020). Histological examination of the tissues 56 days after transplantation revealed growth plate-like structures in all patient-derived clones and their isogenic controls (Figures 5A-5C). Safranin O staining and von Kossa staining showed no difference between the mutants and controls, indicating normal cartilage matrix production and mineralization.

**Figure 5.**
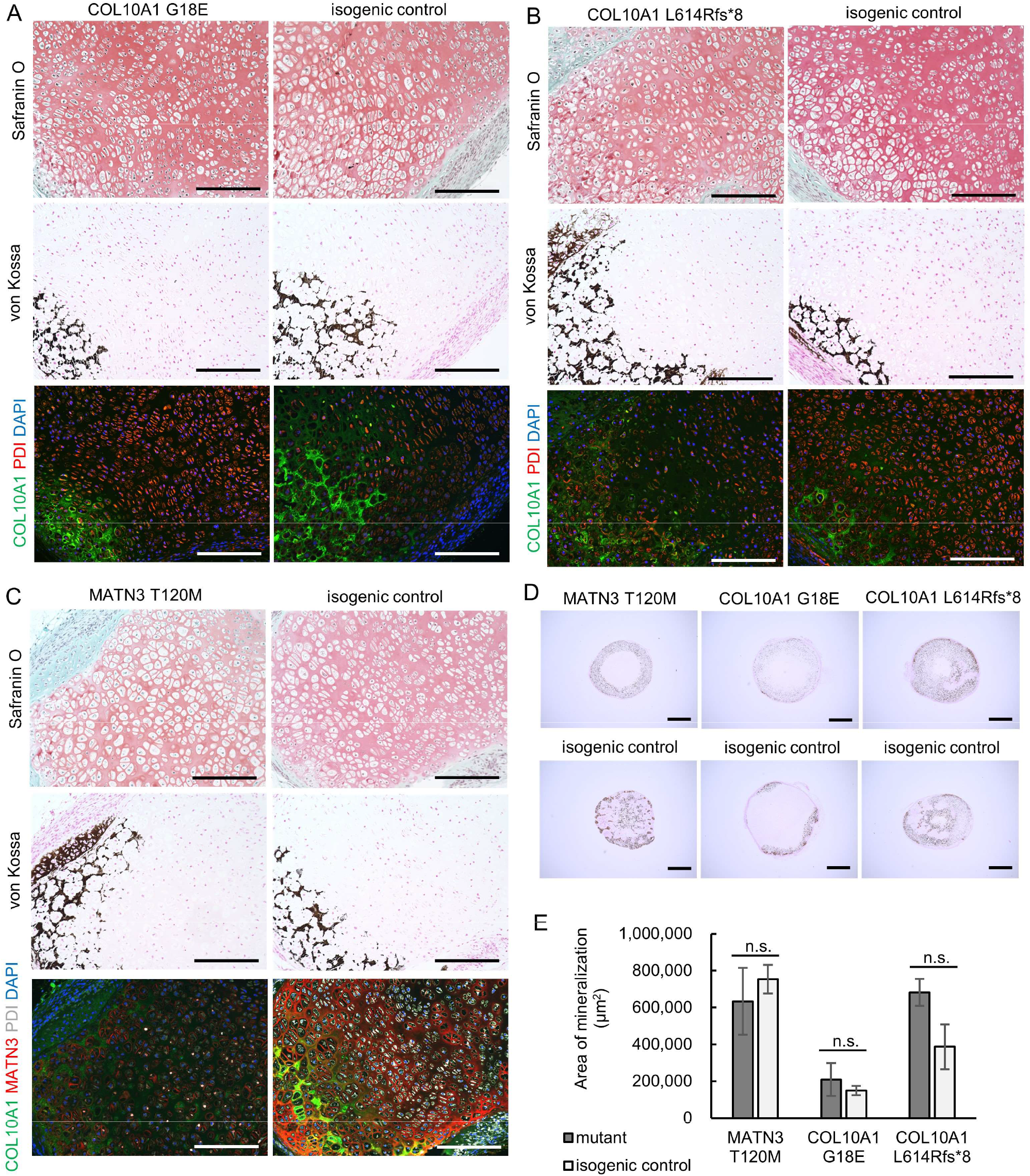
*COL10A1* and *MATN3* mutants show dysregulated hypertrophy without change in mineralization. (A-C) Histology of growth plate-like structures from tissue collected on day 56 after transplantation into immunodeficient mice. (top) Safranin O staining, (middle) von Kossa staining, (bottom) immunostaining. Scale bars, 200 μm. Similar results were observed in two or more growth plate-like structures for each clone. (D) von Kossa staining of mutant (top) or isogenic control (bottom) pellets transplanted on day 28 of HI and collected after 20 days *in vivo*. Similar results were obtained in six pellets transplanted into three different mice (two pellets per mouse). Scale bar, 1 mm. (E) Quantification of von Kossa-positive areas. Values are expressed as mean±SEM. Results are calculated as n=3 from three different mice with two pellets each. (n.s., no significant difference by unpaired two-sided t-test).

However, in *COL10A1* mutants, a near-complete absence of morphologically hypertrophic chondrocytes was found even in areas where the cartilage matrix started to mineralize (Figures 5A and 5B). Columnar chondrocytes were observed, but chondrocyte morphology showed little change throughout the structure and no distinct hypertrophic zone could be observed. This is despite the expression of COL10A1 ahead of the mineralized area, indicating that a lack of hypertrophic morphology does not preclude COL10A1 production and secretion. In the *MATN3* mutant, cells with a hypertrophic morphology were observed throughout the growth plate-like structure, which lacked columnar chondrocytes and a distinct proliferative zone (Figure 5C). MATN3 and COL10A1 were detected weakly throughout the entire structure. Taken together, these results are similar to the morphological abnormalities detected in homozygous model mice (Leighton et al., 2007; Rajpar et al., 2009).

In order to examine whether mineralization is affected as a result of these abnormalities, we transplanted cartilage pellets created using our *in vitro* system into immunodeficient mice on day 28 of HI. Within 20 days, mineralization was detected in all mutants and isogenic controls (Figure 5D), but the area of mineralization was not different between the isogenic pairs (Figure 5E).

### Autophagy inducers and chemical chaperones alleviate ER stress with varying efficacy

We next asked whether our *in vitro* system could be used for drug testing. As autophagy inducers and chemical chaperones have been shown to improve the phenotype in models of skeletal dysplasias (Forouhan et al., 2018; Kawai et al., 2019; Okada et al., 2015; Posey et al., 2014), we assessed their effect on the *COL10A1* and *MATN3* mutants. In the COL10A1 S600P mutant, carbamazepine (CBZ) and rapamycin (RM) decreased the expression of ER stress markers, but rapamycin also led to a decrease of *COL2A1* expression and pellet size, indicating a potential inhibition of cartilage maturation (Figures 6A and 6B). In the MATN3 T120M mutant, trimethylamine N-oxide (TMAO) was most effective in reducing ER stress marker expression without also decreasing *COL2A1* expression or pellet size. TMAO also significantly decreased the ER size in this mutant, while CBZ significantly decreased the ER size in the COL10A1 S600P mutant (Figure 6C). CBZ treatment did, however, not resolve intracellular COL10A1 aggregates, whereas TMAO treatment led to a significant reduction in intracellular retention of MATN3 (Figures 6D and 6E). This indicates that TMAO might help in preventing MATN3 misfolding, but CBZ acts only to reduce the ER stress as a result of COL10A1 accumulation.

**Figure 6.**
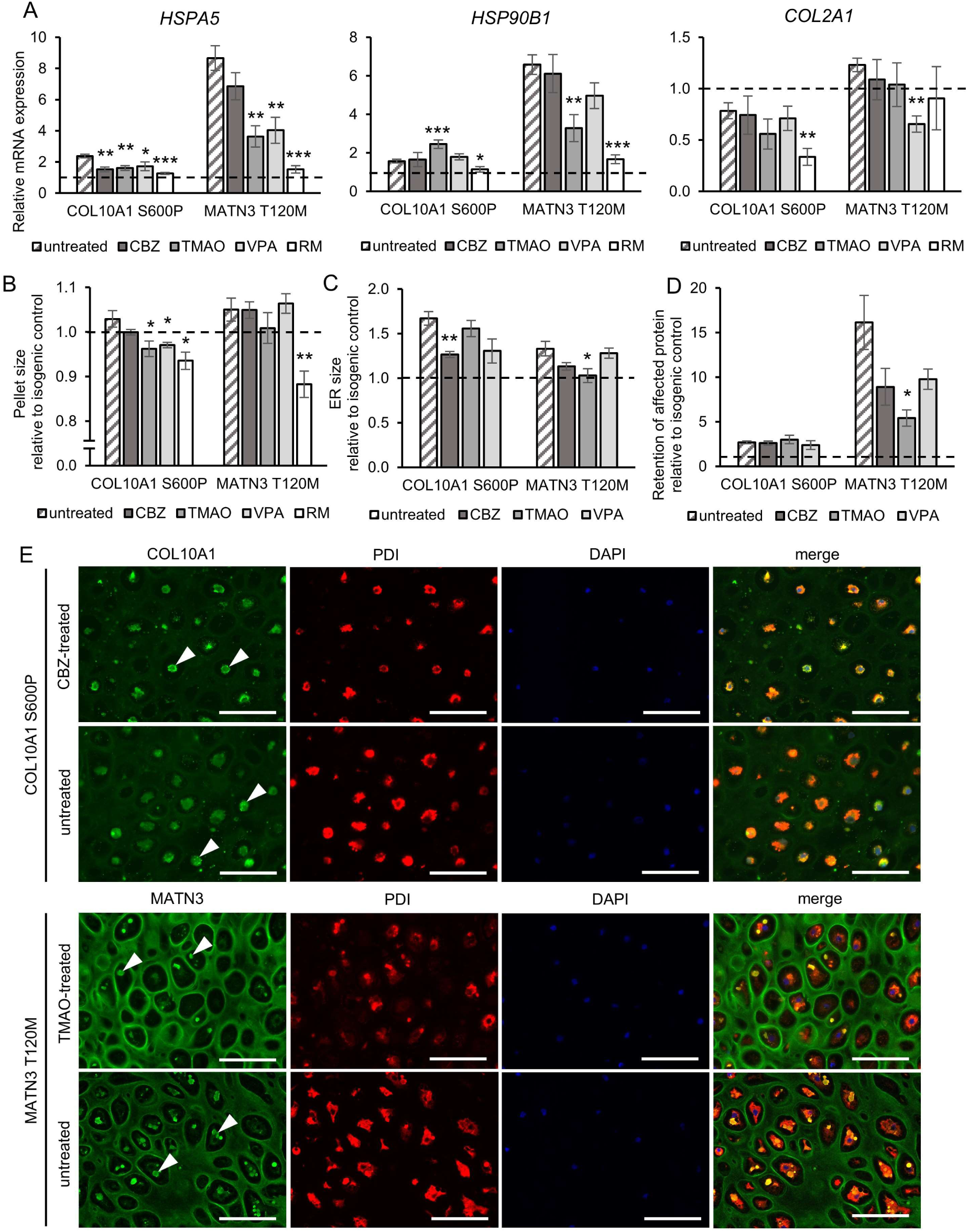
Autophagy inducers and chemical chaperones reduce ER stress in mutants. (A) mRNA expression of ER stress markers and chondrocyte marker *COL2A1* (biologically independent sample number shown in Table S1). (B) Pellet size of treated and untreated mutants. (C) ER size quantified from the area of PDI fluorescence. (D) Intracellular retention quantified from fluorescence intensity of COL10A1 or MATN3 costaining with PDI (intracellular) divided by total fluorescence intensity. (E) Immunostaining of COL10A1 or MATN3 and PDI. Arrowheads indicate intracellular aggregates. Similar results were obtained in four biologically independent experiments. Scale bars, 100 μm. All results are from day 56 of HI and expressed as mean±SEM, relative to the isogenic control (dotted lines). Where not otherwise indicated, samples are n=4 biological replicates. As experiments were performed at the same time, data of untreated mutants and isogenic controls in (A) to (D) are the same as in Figures 2D, 3C, 3D, 4A, 4D, and S5A. CBZ, carbamazepine; TMAO, trimethylamine N-oxide; VPA, valproic acid; RM, rapamycin. (*p < 0.05, **p < 0.01, ***p < 0.001 by unpaired two-sided t-test of treated compared to untreated samples; indication of n.s., no significance, is omitted). See also Figure S6.

The alleviation of ER stress by CBZ and TMAO was not consistently observed in all mutants, however. ER stress markers showed only a tendency or no decrease at all in the COL10A1 G18E, COL10A1 L614Rfs*8, and MATN3 R209P mutants (Figure S6A). Similarly, ER size and intracellular retention showed little or no change in these mutants (Figures S6B-S6E). Taken together, the autophagy inducers and chemical chaperones proved most effective in the mutants that showed the most severe ER stress and intracellular retention, while having less effect on mutants with the milder phenotypes.

### Mutants show common and mutation-specific changes in the transcriptome and metabolome

In order to elucidate the effects of the UPR on various biological processes, we compared the gene expression profile of the mutants and isogenic controls through microarray analysis on day 56 of HI. The number of differentially expressed genes varied depending on the mutation, with the COL10A1 S600P and MATN3 T120M mutants showing the largest number at both the cutoff of p<0.05 and p<0.001 (Figures 7A and 7B). The MATN3 R209P mutant showed the lowest amount of differentially expressed genes, with almost no genes passing the p<0.001 cut-off. Therefore, p<0.05 was used for subsequent analysis.

**Figure 7.**
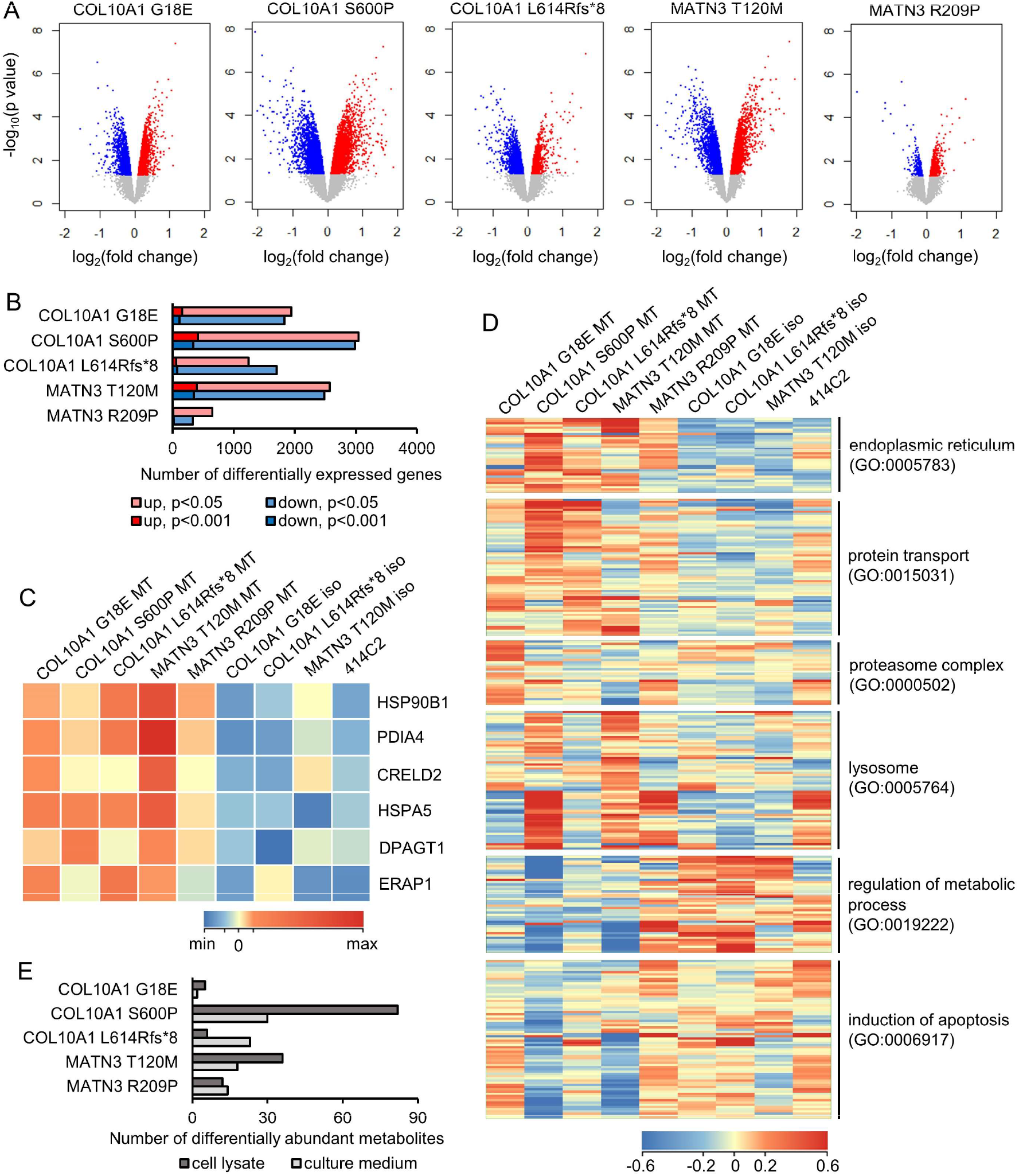
Mutants show both common and mutation-specific transcriptional and metabolic changes. (A) Volcano plot from microarray data of differentially expressed genes (p<0.05 by moderated t-test) of mutants (MT) vs their isogenic controls (iso). Red points, upregulated; blue points, downregulated. (B) Number of differentially expressed genes in each mutant compared to its isogenic control. P value is calculated by moderated t-test. (C, D) Heatmap of genes differentially expressed in every mutant (C) or in at least one mutant (D). The isogenic control of both COL10A1 S600P and MATN3 R209P mutants is denoted as 414C2. Legend indicates level of expression normalized by the median of all samples. See table S2 for the complete list of genes shown in (D) in order. (E) Number of differentially abundant metabolites in each mutant compared to its isogenic control. P value is calculated by two-sided t-test with unequal variances. All results are from n=3 biological replicates from day 56 of HI. See also Figure S7.

Surprisingly, only 81 genes were commonly upregulated in *COL10A1* mutants, and 70 genes were commonly upregulated in *MATN3* mutants (Figure S7A). The number of commonly downregulated genes was 95 in *COL10A1* and 22 in *MATN3* mutants. No overlap of downregulated genes was seen between *COL10A1* and *MATN3* mutants, but 6 genes were upregulated in every mutant. The heatmap of these 6 genes, all of which encode ER-resident proteins, clearly shows the different expression pattern in mutants compared to isogenic controls (Figure 7C). GO analysis of upregulated genes common to either all *COL10A1* or all *MATN3* mutants also showed several ER-related terms (Figure S7B). In contrast, GO terms of commonly downregulated genes included terms related to metabolic processes and proteoglycan synthesis.

Looking at genes differentially expressed in at least one, but not necessarily all, of the mutants, we found transcriptomic changes that were common to most mutants as well as some that were seen in only one mutant (Figure 7D; complete list in Table S2). Genes related to protein transport, such as coatomer subunits, vesicular trafficking proteins, and cargo loading proteins, tended to be upregulated in mutants. Surprisingly, genes encoding proteins of the proteasome complex were highly upregulated only in the COL10A1 G18E mutant, whereas lysosomal proteins were upregulated only the COL10A1 S600P and MATN3 T120M mutants, indicating that different degradation pathways are activated depending on the mutation. Genes related to the regulation of metabolic processes were downregulated in all mutants except MATN3 R209P. The COL10A1 S600P and MATN3 T120M mutants again showed a similar pattern of expression in genes related to the induction of apoptosis, which were downregulated in only these two mutants.

Since metabolic process regulation was affected in mutants, and metabolites can be useful biomarkers of disease, we performed gas chromatography-mass spectrometry (GC-MS) using both cell lysates and culture medium from day 56 of HI for metabolome analysis. The COL10A1 S600P and MATN3 T120M mutants, which had shown the largest number of differentially expressed genes, also showed the largest amount of differentially abundant metabolites (Figure 7E; complete list in Tables S3 and S4). In the COL10A1 S600P mutant, both the culture medium and the cell lysate showed a depletion of some metabolites such as glutamic and aspartic acid, and arginine biosynthesis was also identified as an affected pathway in both medium and lysate (Figure S7C). In fact, arginine biosynthesis was affected in the lysate of all *COL10A1* mutants, although no common differentially abundant metabolites were found. *MATN3* mutants showed neither common pathways nor metabolites, indicating that metabolic changes were mostly mutation-specific.

## Discussion

In this study, we developed a robust, serum-free protocol to induce hypertrophic chondrocytes from iPSCs to model chondrodysplasias in a 3D culture system. This protocol mimics the process of differentiation in growth plate chondrocytes, with cells characteristic of hypertrophic chondrocytes in their morphology as well as their gene expression being obtained after 56 days in 3D culture. The appearance of hypertrophic chondrocytes from the periphery of the pellet on day 28, approaching and then reaching the center of the pellet by day 56, indicates the active promotion of hypertrophic differentiation by factors diffusing into the pellet from the medium. In addition, genes that are highly upregulated in the physis but not articular cartilage, such as *MATN3* and *COL9A2* (Paradise et al., 2018), were expressed in the course of induction, supporting the validity of our model to research disorders of the growth plate.

Using this model, we could confirm the presence of intracellular aggregates and ER stress in human *COL10A1* and *MATN3* mutants *in vitro*. However, changes in cell morphology and chondrocyte maturation markers were small in this system. Only *IHH*, which is part of the IHH-PTHrP feedback loop that regulates chondrocyte differentiation in the growth plate (Kobayashi et al., 2002), showed lower expression in mutants, whereas morphological abnormalities required an *in vivo* environment to appear. Together with the expression of *COL2A1* remaining high on day 56, this suggests that our *in vitro* system may still lack the proper regulatory network, which may require a zonal arrangement that exists only *in vivo*. Therefore, an *in vitro* system that includes both proliferating and hypertrophic chondrocytes in separate zones, similar to a growth plate *in vivo*, may greatly expand the capabilities of chondrodysplasia-specific iPSCs to precisely recapitulate the disease phenotype.

One of the greatest barriers to understanding chondrodysplasias is their heterogeneity among patients with mutations in the same disease-causing gene. This heterogeneity may originate from the distinct genetic backgrounds of each patient, as well as from differences in the type and location of their mutation. Our *in vitro* system has allowed us to model two different chondrodysplasias at once, with two or three different mutations each, with isogenic controls for each mutation, making it possible to precisely evaluate the impact of each mutation on the phenotype. Using this system, we found that the COL10A1 S600P and MATN3 T120M mutations generally resulted in the most severe ER stress, highest retention of the affected protein, and largest transcriptomic changes. The patient with the MATN3 T120M mutation had a short stature (−2.2 SD) and a severe phenotype in the radiological findings, with skeletal abnormalities reminiscent of spondyloepimetaphyseal dysplasia, matrilin 3 type (SEMD-MATN3; OMIM #608728), which is caused by homozygous *MATN3* mutations (Borochowitz et al., 2004). Since the SNP rs187943382, causing MATN3 V220A, was found in the healthy allele of *MATN3* in this patient, we cannot exclude that it could exacerbate the pathology caused by T120M, as both are within the vWFa domain and may act in synergy to worsen the misfolding. Because V220A has also been detected in healthy controls (Kim et al., 2011), and the rescue of the MATN3 T120M mutant showed no intracellular accumulation of MATN3, we propose that this SNP may only worsen existing MED pathology without causing it on its own. In contrast to the MATN3 T120M mutant, the MATN3 R209P mutant showed only very mild ER stress, intracellular retention, and transcriptomic changes. The height of the R209P patient was in the 51st percentile (Kim et al., 2011), indicating that this mutation may potentially result in fewer structural changes in MATN3 and a milder phenotype than other mutations, which caused very short statures.

However, the MCDS patients all showed similar symptoms typical of this chondrodysplasia. The difference was the type of the *COL10A1* mutation, with G18E causing an amino acid substitution in the signal peptide, S600P causing an amino acid substitution in the NC1 domain, and L614Rfs*8 being a frameshift mutation causing an early stop codon in the NC1 domain. Since the NC1 domain is critical for the trimerization of COL10A1 chains (Bogin et al., 2002), both mutations in this domain are expected to cause misfolding of the protein, resulting in ER stress. As we observed no NMD resulting from the L614Rfs*8 mutation, this frameshift mutation likely acts in a gain-of-function manner similar to S600P, rather than leading to degradation and haploinsuffiency. The G18E mutation, however, likely leads to impaired signal peptide cleavage (Chan et al., 2001), so that the COL10A1 chain cannot be released into the ER lumen in a timely manner after translation. While it may associate with other COL10A1 chains, we found that the secretion into the extracellular matrix (ECM) was significantly reduced, so that it likely accumulates within the ER and triggers the UPR, resulting in a phenotype similar to that caused by mutations in the NC1 domain. Interestingly, the G18E mutant was the only mutant with genes of the proteasomal complex being highly upregulated, suggesting that this mutation may uniquely promote the export of the mutant COL10A1 from the ER. Further evidence of proteasomal degradation of COL10A1 G18E mutant chains is provided in a previous report showing degradation of these chains was inhibited by proteasome inhibitors (Chan et al., 2001).

Whether apoptosis is a disease mechanism in mild chondrodysplasias such as MCDS is controversial (Cameron et al., 2015). Generally, severe and prolonged ER stress promotes the apoptotic pathway, while mild, short-term ER stress leads to an adaptive response in the cell (Hetz, 2012). In our *in vitro* system, we observed no increase of apoptosis in mutant hypertrophic chondrocytes; on the contrary, some mutants showed a lower amount of apoptosis compared to their isogenic controls in the TUNEL assay. These findings were confirmed in our transcriptome analysis, where genes involved in the induction of apoptosis, such as *BAX*, *RIPK1*, *FAS*, and *DAPK1*, were significantly downregulated in the COL10A1 S600P and MATN3 T120M mutants. These mutants had the highest ER stress, indicating that ER stress caused by *COL10A1* and *MATN3* mutations is not severe enough to induce the apoptotic pathway and may instead trigger the adaptive response to inhibit cell death. The other mutants did not show such reduced expression of apoptosis-related genes, suggesting that the ER stress may have been mild enough not to trigger the full adaptive response. A similar negative regulation of apoptosis has also been reported in the hypertrophic zone of an MCDS model mouse (Wang et al., 2018), but contrary to this model, *FGF21* or *ATF4* expression was not increased in our *COL10A1* mutants.

Gene expression profiling showed the upregulation of ER-related genes as a common GO term, but only six genes were commonly upregulated in all mutants. All six encode ER-resident proteins, including ER stress markers *HSP90B1, CRELD2*, and *HSPA5*, whose expression we also found to be significantly increased in most mutants by qPCR. The other three genes were *PDIA4, DPAGT1*, and *ERAP1*, all of which have also been implicated in ER stress or the UPR (Ferrari and Söling, 1999; Heifetz et al., 1979; Thomaidou et al., 2020). On the other hand, GO terms of downregulated genes were related to the synthesis of proteoglycan, a major component of the ECM, suggesting that ER stress may lead to a deterioration of ECM quality and interfere with the columnar arrangement of the growth plate.

Autophagy inducers and chemical chaperones, which are expected to reduce ER stress by respectively stimulating partial digestion of the ER or assisting the correct folding of the protein, did not consistently ameliorate the phenotype in all *COL10A1* and *MATN3* mutants. Therefore, drugs targeting a different pathway or mechanism may be required for effective treatment of chondrodysplasias with specific mutations, even if such pathways are not affected in every mutant. One group of genes upregulated in most, but not all, of the mutants involved the protein transport. These genes encoded proteins including the coatomer subunits COPB2, COPE, COPG, and COPZ1; coat-recruiting proteins such as ARF3 and ARF4; COPII-binding protein subunits SEC22B and SEC23IP; KDEL receptors KDEL1, KDEL2 and KDEL3; and collagen-packaging protein MIA3. As a similar increase of protein transport-related gene expression in the UPR has been previously reported (Murray et al., 2004), the upregulation of these genes is likely part of the UPR elicited by the intracellular accumulation of COL10A1 and MATN3. Therefore, supporting the export of proteins, regardless of their misfolded conformation, may be a promising avenue to explore as treatments for chondrodysplasias with intracellular protein retention where suppressing ER stress alone may not be sufficient.

In order to find suitable biomarkers for drug screening, we examined the metabolome for any changes that could be targetable with drugs. The only pathway commonly affected in all *COL10A1* mutants was arginine biosynthesis, with ornithine being significantly more abundant in the lysates of G18E and L614Rfs*8 mutants, and arginine, fumarate, glutamate, aspartate, and 2-ketoglutarate being significantly less abundant in the S600P mutant compared to the isogenic control. We found mild differential expression of arginine metabolism-related genes such as *ARG2, NOS1, NOS2*, and *ASS1* in one or more *COL10A1* mutants in the transcriptome, but their up-or downregulation was not consistent across different mutants, indicating that changes in this pathway do not occur universally from *COL10A1* mutations. Given that arginine depletion has been shown to trigger the UPR (García-Navas et al., 2012), we hypothesize that changes in arginine metabolism in *COL10A1* mutants could be an adaptive response to the ER stress. If this is the case, modulation of arginine metabolism may help alleviate the phenotype in mutants with insufficient response to other ER stress suppressors.

Taken together, our results reveal how chondrodysplasia-causing mutations, despite the common activation of the UPR, manifest in differently affected pathways and varying degrees of phenotype severity. Such differences would be missed if only one particular mutation or model mouse was examined. Therefore, iPSC-based models will certainly become a valuable tool to properly assess a multitude of rare patient mutations, shed light on the genotype-phenotype relationship, and perform high throughput drug screening. Since this system recapitulates every stage of chondrocyte differentiation in the growth plate, we expect it to be fully applicable to most other chondrodysplasias, as well, which are individually rare diseases, but taken together cause significant morbidity in large populations.

## Experimental procedures

### Establishment of iPSC lines and isogenic controls

Patient iPSC lines were established from skin fibroblasts (MCDS patient #1) or peripheral blood mononuclear cells (MCDS patient #2 and MED patient) as previously described (Okita et al., 2011). Gene-corrected rescues were created using the CRISPR/Cas9 system. Further details are available in the Supplemental Information, including guide RNAs and repair templates in Table S5. All experiments with human subjects were performed with written informed consent and approved by the Ethics Committee of the Department of Medicine and Graduate School of Medicine, Kyoto University, the Ethics Committee of the Shiga Medical Center for Children, and the Ethical Committee of RIKEN Yokohama Institute.

### Sclerotome and hypertrophic induction

All iPSCs were maintained feeder-free on dishes coated with iMatrix-511 silk (Nippi) in StemFit AK02N (Ajinomoto) with 50 U penicillin and 50 μg/ml streptomycin (Gibco). Cells were passaged at a density of 1.1 × 10^3^ to 3.2 × 10^3^ cells/cm^2^ five days before induction. SI was performed as previously described (Matsuda et al., 2020). On day 6 of SI, cells were detached and resuspended in CDMi base medium containing 100 nM SAG (Calbio), 600 nM LDN193189 (Stemgent), and 10 μM Y-27632 (Wako). Cells were seeded into low attachment 96 well plates (SUMILON) at 2.5 × 10^5^ cells/well.

On the next day, i.e., day 0 of HI, the medium was changed to HI base medium supplemented with 40 ng/ml PDGF-BB (R&D) and 0.1 μM dexamethasone (Wako). From day 6 of HI, 10 ng/ml TGFβ3 (R&D) was additionally supplemented. From day 10 of HI, PDGF-BB was removed and 50 ng/ml BMP4 (R&D) was added. From day 14, dexamethasone and TGFβ3 were removed, and 10 nM triiodothyronine (T3) (Sigma) was added. From day 28, 10 mM β-glycerophosphate (Sigma) was added. For the drug-treated groups, 20 μM carbamazepine (CBZ) (Sigma), 50 mM trimethylamine N-oxide (TMAO) (Sigma), 200 μM valproic acid (VPA) (Sigma), or 10 nM rapamycin (RM) (MedChem Express) was added from day 28 to 56 of HI. The CDMi and HI base medium compositions and other information about induction are detailed in the Supplemental Information, including key reagents in Table S7.

### Animal experiments

To observe growth plate-like structures, cells were detached on day 6 of SI and resuspended in CDMi containing 100 nM SAG, 600 nM LDN193189, and 10 μM Y-27632 at a concentration of up to 1 × 10^8^ cells/ml. This cell suspension was mixed 1:1 with matrigel (BD) and 100 μl were injected subcutaneously into a minimum of six male immunodeficient NOD/ShiJic-scid Jcl (NOD-SCID) mice (CLEA Japan). The transplanted tissue was collected after 56 days. To quantify mineralization of cartilage, pellets were transplanted subcutaneously into three male NOD-SCID mice on day 28 of HI. Pellets were collected after 20 days. All animal experiments were approved by the UCSF Institutional Animal Care and Use Committee and performed in accordance with the Regulations on Animal Experimentation at Kyoto University.

### Expression analysis and flow cytometry

After RNA extraction using the RNeasy Micro or Mini Kit (QIAGEN), quantitative PCR (qPCR) was performed with the Thunderbird SYBR qPCR Mix (Toyobo). Microarray analysis was performed with the Human Gene 1.0ST Array (Affymetrix). Protein expression was analyzed by Simple Western (ProteinSimple). Flow cytometry was performed using the FACS Aria II (BD) as previously described (Matsuda et al., 2020) with minor modifications. More details are in the Supplemental Information, including primer information in Table S6. The microarray data are available in the GEO database under the accession number GSE148728.

### Histological analysis

*In vitro* pellets and *in vivo* tissue were fixed in 4% paraformaldehyde for 2 days before paraffin embedding. Staining protocols are detailed in the Supplemental Information.

### Gas chromatography coupled to mass spectrometry (GC-MS)

For metabolome analysis, GC-MS was performed using the GCMS-TQ8030 (Shimadzu) with culture medium or pellet extracts prepared on day 56 of HI. Data were acquired and peaks were processed with the GCMS solution software (Shimadzu). Additional protocol details can be found in the Supplemental Information.

## Author Contributions

Y.P., S.K., and J.T. designed the study. Y.P. developed the hypertrophic induction protocol, performed the gene editing and disease modeling, analyzed all experiments, and drafted the manuscript. M.W. performed the GC-MS and analyzed the metabolomics data. S.N. and M.N. assisted in the *in vitro* and *in vivo* experiments, respectively. S.T. supervised the establishment of patient iPSC lines. C.A. devised the sclerotome induction and transplantation protocols. J.-Y.X. and Z.W. performed the mutation search by NGS. K.F., M.T., T.F., and S.I. provided patient samples for the establishment of iPSC lines. J.T. supervised the study and revised the manuscript, which was further reviewed by all authors.

## Acknowledgements

We thank J. Ma and T. Nakashima for their assistance in the *in vitro* and *in vivo* experiments, respectively; H. Yoshitomi, Y. Jin, and T. Takarada for comments and discussion; and Y. Yamanaka for advice on sclerotome differentiation and transplantation. Preparation of tissue slides and staining was supported by the Center for Anatomical, Pathological and Forensic Medical Research, Graduate School of Medicine, Kyoto University, and Applied Medical Research Laboratory. Karyotyping was performed by chromocenter, Inc. This study was supported by Grant-in-Aid for the Acceleration Program for Intractable Disease Research Utilizing Disease Specific iPS Cells (AMED) for S.I. and J.T.

