## Supplementary Information for "Human iPSC-derived hypertrophic chondrocytes reveal a mutation-specific unfolded protein response in chondrodysplasias"

**Inventory of Supplemental Information**

- Supplemental Figures S1-S7
- Supplemental Tables S1-S7
- Supplemental Experimental Procedures
- Supplemental References

**Supplemental Figures**

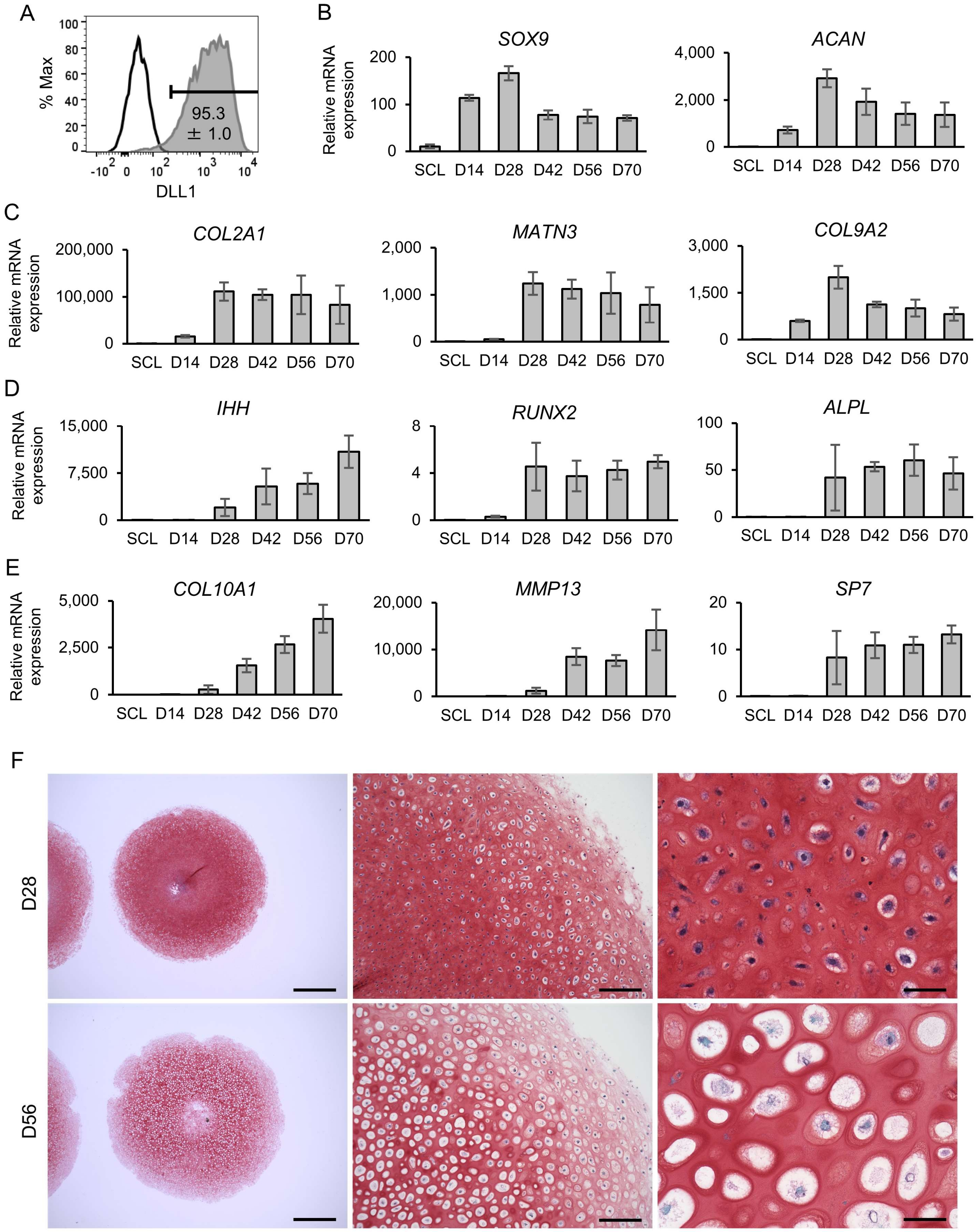

**Figure S1. Differentiation of hypertrophic chondrocytes from 1231A3 iPSCs. Related to Figure 1**

(A) Representative result of DLL1-positive cells (compared to isotype control) on day 2 of sclerotome induction (SI) with the mean and SEM (standard error of the mean) of four biological replicates displayed.

(B-E) Expression of early (B), proliferating (C), pre-hypertrophic (D) and hypertrophic (E) chondrocyte markers over time from sclerotome (SCL) on day -1 to day 70 of hypertrophic induction (HI). Values are relative to the chondroblastic osteosarcoma cell line ANOS, which stably expresses these markers, and are shown as mean±SEM (n=4 from biologically independent experiments).

(F) Representative result of Safranin O staining of pellets from day 28 and 56 of HI. Similar results were obtained in three biologically independent experiments. Scale bars, (left) 1 mm, (middle) 200 μm, (right) 50 μm.

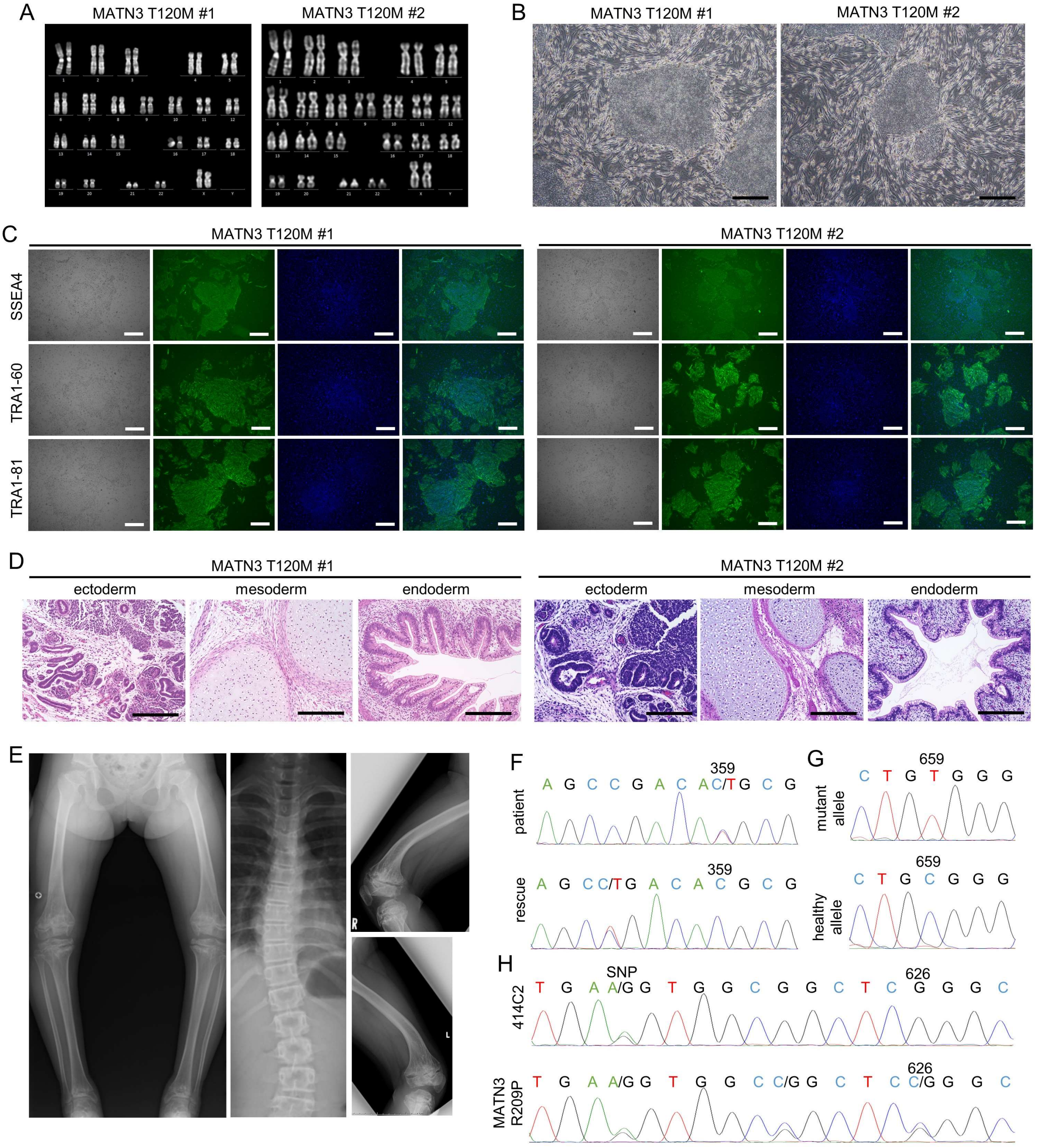
A
MATN3 T120M #1
MATN3 T120M #2
B
MATN3 T120M #1
MATN3 T120M #2
C
MATN3 T120M #1
MATN3 T120M #2
SSEA4
TRA1-60
TRA1-81
D
MATN3 T120M #1
MATN3 T120M #2
ectoderm
mesoderm
endoderm
ectoderm
mesoderm
endoderm
E
F
patient
A G C C G A C A C/T G C G
359
rescue
A G C C/T G A C A C G C G
359
G
mutant allele
C T G T G G G
659
healthy allele
C T G C G G G
659
H
414C2
SNP
T G A A/G G T G G C G G C T C G G G C
626
MATN3 R209P
T G A A/G G T G G C C/G G C T C C/G G G C
626

**Figure S2. Multiple epiphyseal dysplasia (MED) patient phenotype, validation of iPSC clones and sequence of an additional *MATN3* mutant clone. Related to Figure 2**

(A) Karyotype analysis of MED patient-derived clones #1 and #2, showing a normal karyotype of 46,XX.

(B) Phase-contrast image of iPSC clones. Scale bars, 100 µm.

(C) Brightfield image (left) and immunostaining (right) of each clone. Pluripotent markers are shown in green with the nuclear counterstain DAPI in blue. Scale bars, 100 µm.

(D) Tissue formed from teratomas of each clone, showing differentiation into ectoderm, mesoderm and endoderm. Scale bars, 200 µm.

(E) Patient x-ray image of the lower limbs (left), spine (middle), right femur (upper right) and left femur (lower right).

(F) Sequence of the patient showing the heterozygous *MATN3* c.359C>T (p.T120M) mutation (top) and the gene-corrected rescue (bottom).

(G) Single allele sequence of the mutant allele (top) and healthy allele (bottom), showing the healthy allele containing the SNP rs187943382, *MATN3* c.659T>C (p.V220A).

(H) Sequence of the wild type iPSC line 414C2 (top), showing the SNP in *MATN3* used for allele-specific targeting, and the mutant 414C2 with the heterozygous *MATN3* c.626G>C (p.R209P) mutation (bottom).

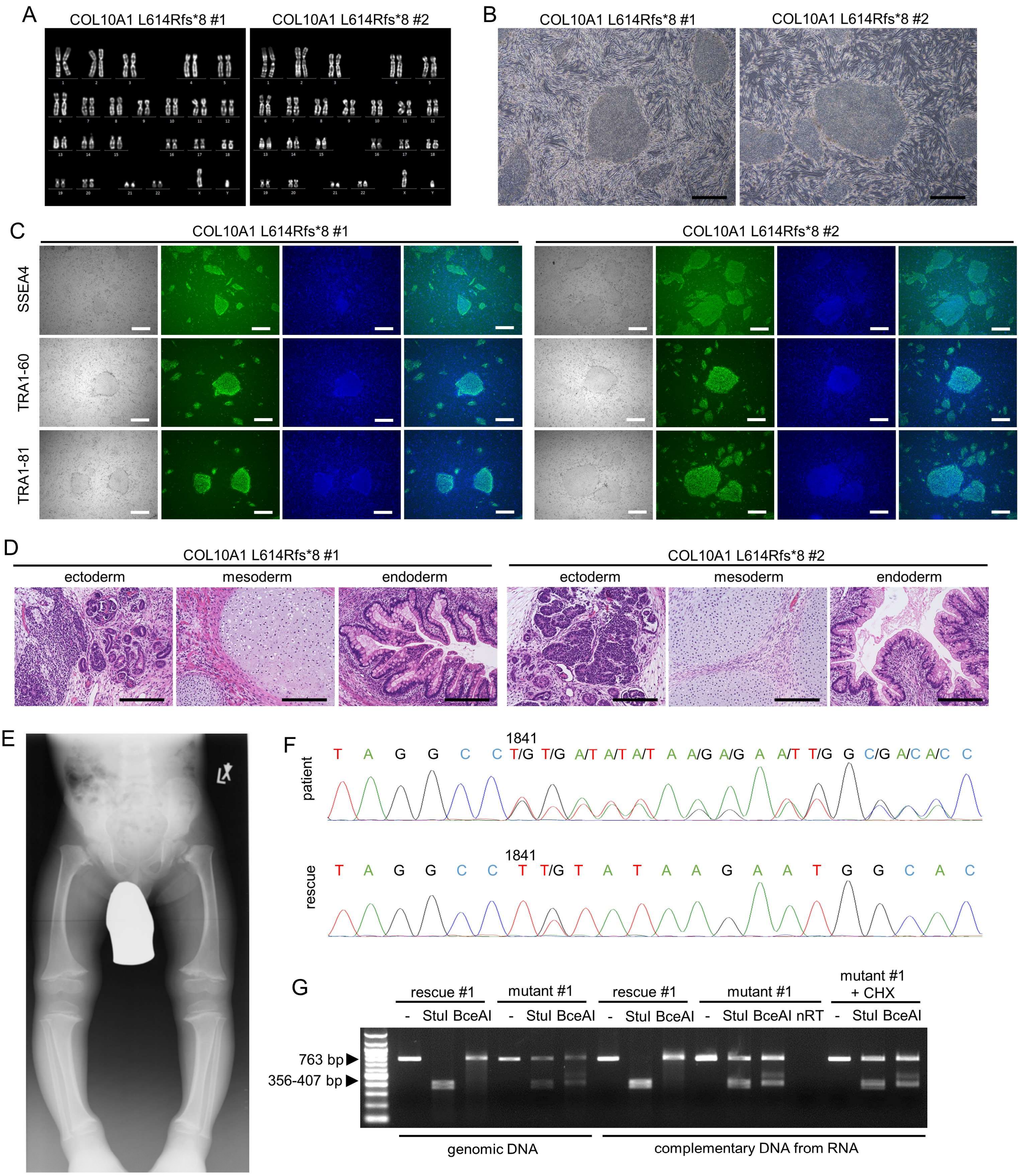
A
COL10A1 L614Rfs*8 #1
COL10A1 L614Rfs*8 #2
B
COL10A1 L614Rfs*8 #1
COL10A1 L614Rfs*8 #2
C
COL10A1 L614Rfs*8 #1
COL10A1 L614Rfs*8 #2
SSEA4
TRA1-60
TRA1-81
D
COL10A1 L614Rfs*8 #1
COL10A1 L614Rfs*8 #2
ectoderm
mesoderm
endoderm
ectoderm
mesoderm
endoderm
E
F
1841
patient
T A G G C C T/G T/G A/T A/T A/T A A/G A/G A A/T T/G G C/G A/C A/C C
1841
rescue
T A G G C C T T/G T A T A A G A A T G G C A C
G
rescue #1
mutant #1
rescue #1
mutant #1
mutant #1 + CHX
- StuI BceAI
- StuI BceAI
- StuI BceAI
- StuI BceAI nRT
- StuI BceAI
763 bp
356-407 bp
genomic DNA
complementary DNA from RNA

**Figure S3. Metaphyseal chondrodysplasia type Schmid (MCDS) patient #1 phenotype and validation of iPSC clones. Related to Figure 2**

(A) Karyotype analysis of clones #1 and #2, showing a normal karyotype of 46,XY.

(B) Phase-contrast image of iPSC clones. Scale bars, 100 µm.

(C) Brightfield image (left) and immunostaining (right) of each clone. Pluripotent markers are shown in green with the nuclear counterstain DAPI in blue. Scale bars, 100 µm.

(D) Tissue formed from teratomas of each clone, showing differentiation into ectoderm, mesoderm and endoderm. Scale bars, 200 µm.

(E) Patient x-ray image of the lower limbs.

(F) Sequence of the patient showing the heterozygous *COL10A1* c.1841_1841delT (p.L614Rfs*8) mutation (top) and the gene-corrected rescue (bottom).

(G) Electrophoresis after amplifying the *COL10A1* region containing the mutation by PCR and processing the amplicons by restriction enzyme StuI (recognizing only the wild type allele) or BceAI (recognizing only the mutant allele). The mutant allele is clearly visible and not changed by NMD (nonsense-mediated decay) inhibitor CHX (cycloheximide), showing that the early stop codon in L614Rfs*8 does not lead to NMD. The nRT (no reverse transcriptase) negative control shows no band, indicating that no DNA contamination is present. RNA was taken from samples on day 56 of HI. Similar results were obtained in three biologically independent experiments.

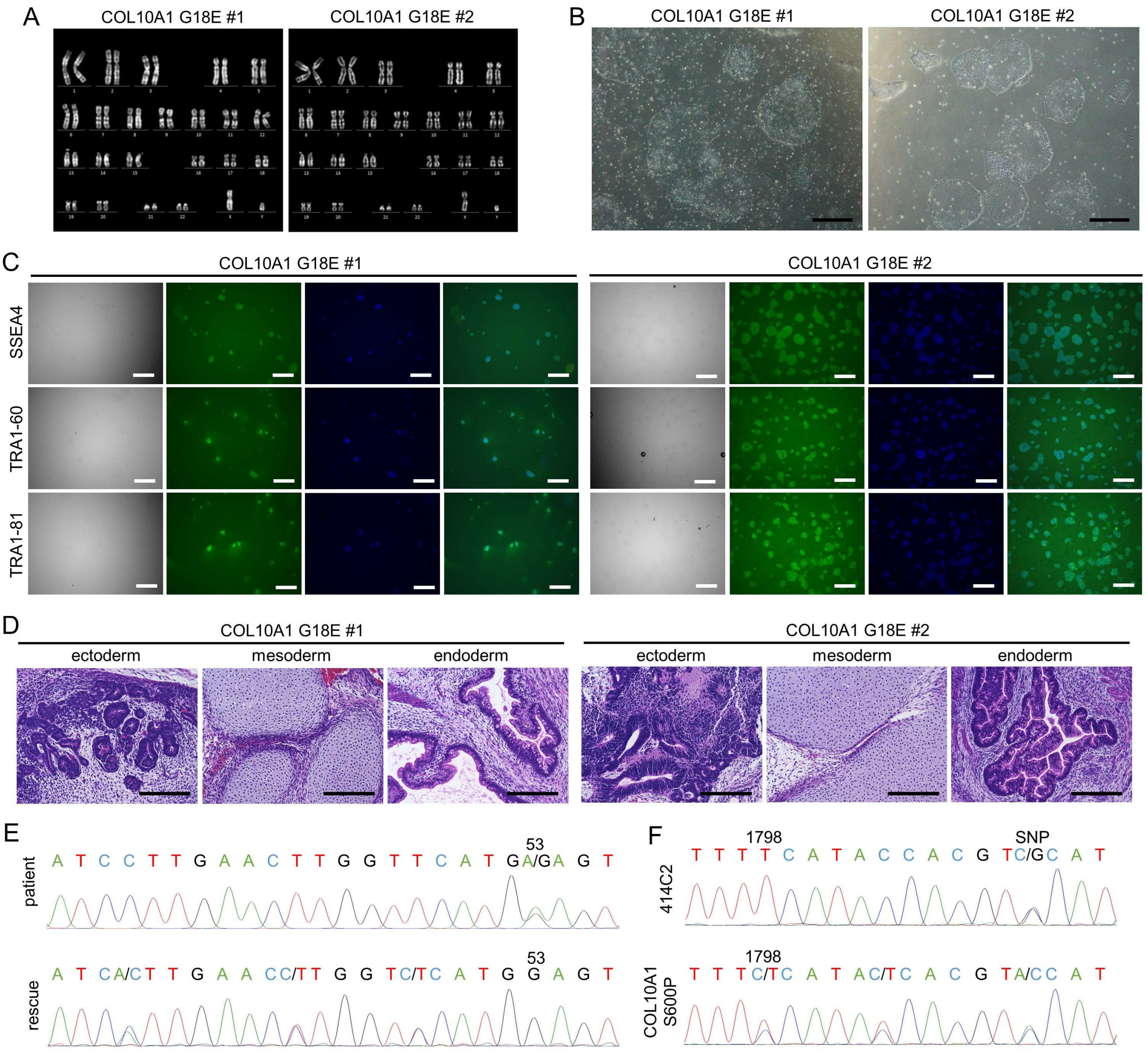

A
COL10A1 G18E #1
COL10A1 G18E #2
B
COL10A1 G18E #1
COL10A1 G18E #2
C
COL10A1 G18E #1
COL10A1 G18E #2
SSEA4
TRA1-60
TRA1-81
D
COL10A1 G18E #1
COL10A1 G18E #2
ectoderm
mesoderm
endoderm
E
patient
53
A T C C T T G A A C T T G G T T C A T G A/G A G T
rescue
53
A T C A/C T T G A A C C/T T G G T C/T C A T G G A G T
F
414C2
1798
SNP
T T T T C A T A C C A C G T C/G C A T
COL10A1 S600P
1798
T T T C/T C A T A C/T C A C G T A/C C A T

**Figure S4. Validation of iPSC clones from MCDS patient #2 and sequence of an additional *COL10A1* mutant clone. Related to Figure 2**

(A) Karyotype analysis of clones #1 and #2, showing a normal karyotype of 46,XY.

(B) Phase-contrast image of iPSC clones. Scale bars, 100 µm.

(C) Brightfield image (left) and immunostaining (right) of each clone. Pluripotent markers are shown in green with the nuclear counterstain DAPI in blue. Scale bars, 100 µm.

(D) Tissue formed from teratomas of each clone, showing differentiation into ectoderm, mesoderm and endoderm. Scale bars, 200 µm.

(E) Sequence of the patient showing the heterozygous *COL10A1* c.53G>A (p.G18E) mutation (top) and the gene-corrected rescue (bottom).

(F) Sequence of the wild type iPSC line 414C2 (top), showing the SNP in *COL10A1* used for allele-specific targeting, and the mutant 414C2 with the heterozygous *COL10A1* c.1798T>C (p.S600P) mutation (bottom).

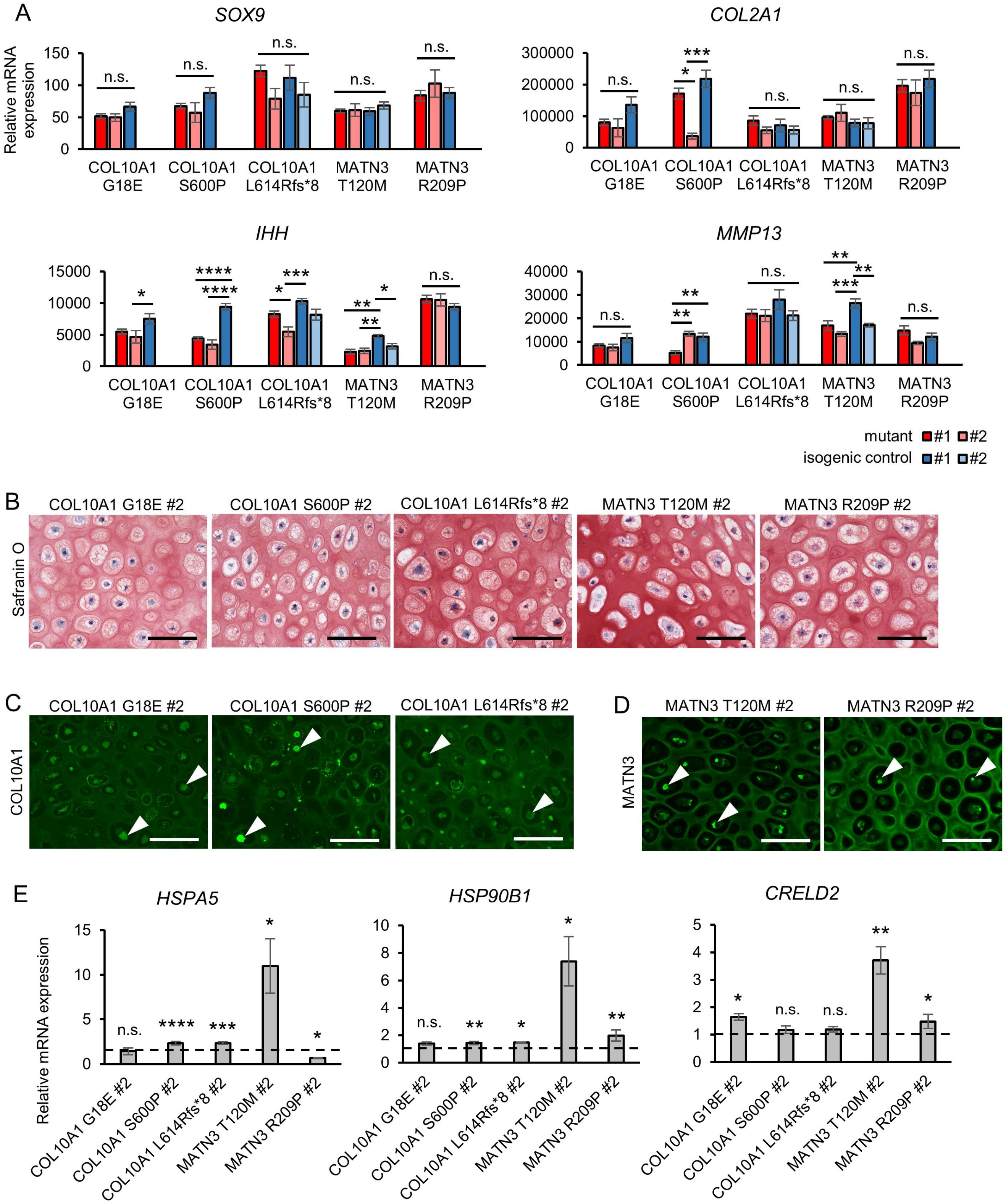
A
SOX9
COL2A1
IHH
MMP13
Relative mRNA expression
n.s.
*
**
***
****
COL10A1 G18E
COL10A1 S600P
COL10A1 L614Rfs*8
MATN3 T120M
MATN3 R209P
mutant #1 #2
isogenic control #1 #2
B
COL10A1 G18E #2
COL10A1 S600P #2
COL10A1 L614Rfs*8 #2
MATN3 T120M #2
MATN3 R209P #2
Safranin O
C
COL10A1
D
MATN3
E
HSPA5
HSP90B1
CRELD2

**Figure S5. Additional mutant clones also show intracellular retention and ER stress without disruption of chondrogenic differentiation. Related to Figures 2, 3, and 4**

(A) mRNA expression of chondrocyte markers of various stages. Values are relative to ANOS (biologically independent sample number shown in Table S1). Statistical analysis by ANOVA with post-hoc Tukey HSD.

(B) Representative result of Safranin O staining. Similar results were obtained in three biologically independent experiments. Scale bars, 100 µm.

(C, D) Immunostaining of COL10A1 (C) or MATN3 (D). Similar results were obtained in three biologically independent experiments. Arrowheads indicate intracellular aggregates. Scale bars, 100 µm.

(E) mRNA expression of ER stress markers. Values are relative to the respective isogenic control (biologically independent sample number shown in Table S1). Dotted lines indicate the value=1 of the isogenic controls. Statistical analysis by unpaired two-sided t-test.

All results are from day 56 of HI and expressed as the mean±SEM. (n.s. no significant difference, $^{*}p < 0.05$, $^{**}p < 0.01$, $^{***}p < 0.001$, $^{****}p < 0.0001$).

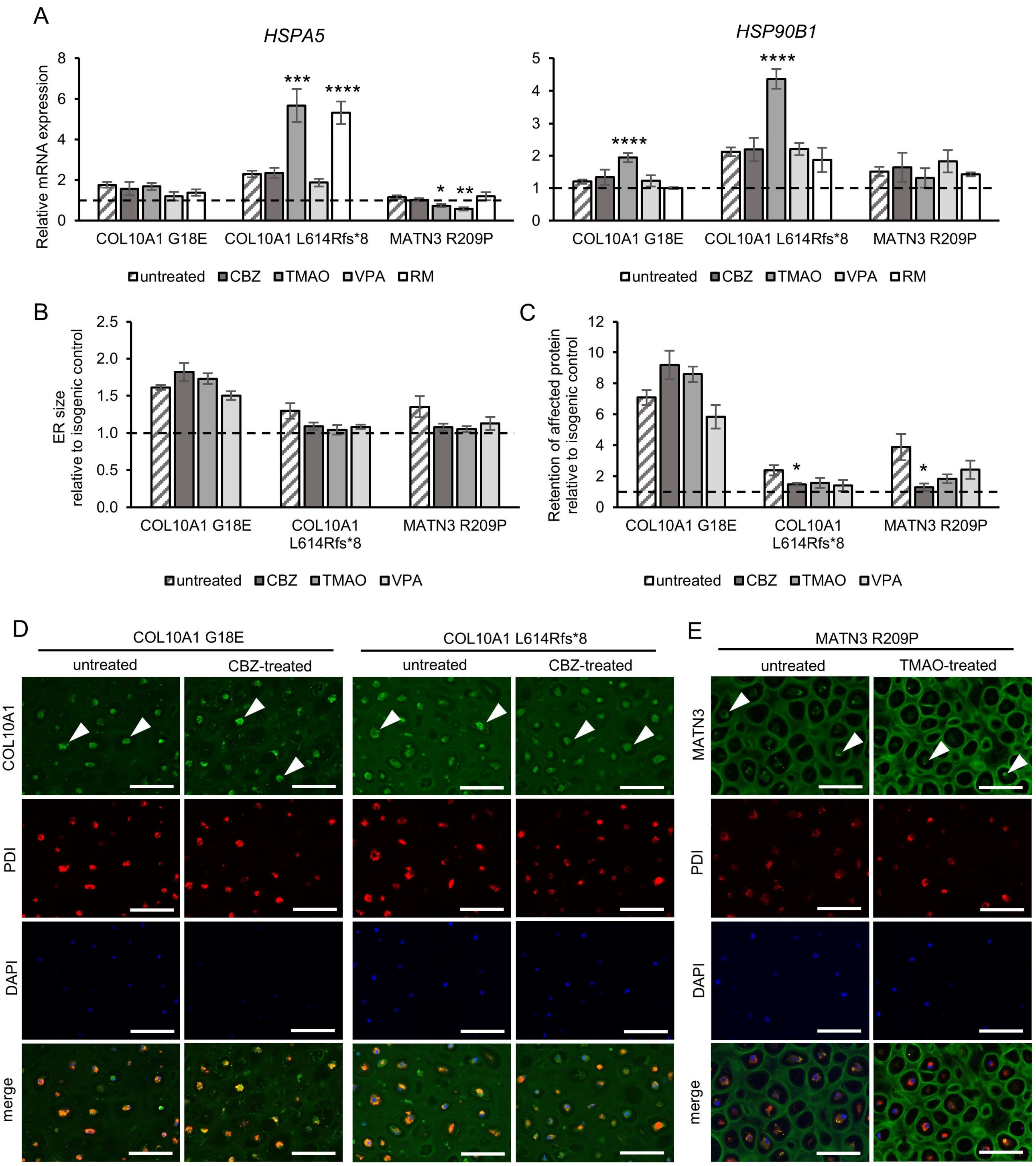
A
HSPA5
HSP90B1
Relative mRNA expression
COL10A1 G18E
COL10A1 L614Rfs*8
MATN3 R209P
untreated
CBZ
TMAO
VPA
RM
B
ER size relative to isogenic control
C
Retention of affected protein relative to isogenic control
D
E
CBZ-treated
TMAO-treated
COL10A1
MATN3
PDI
DAPI
merge

**Figure S6. Drug efficacy varies depending on the mutation. Related to Figure 6**

(A) mRNA expression of ER stress markers (biologically independent sample number shown in Table S1).

(B) ER size quantified from the area of PDI fluorescence.

(C) Intracellular retention quantified from fluorescence intensity of COL10A1 or MATN3 co-staining with PDI (intracellular) divided by total fluorescence intensity.

(D, E) Immunostaining of COL10A1 (D) or MATN3 (E) and PDI. Arrowheads indicate intracellular aggregates. Similar results were obtained in four biologically independent experiments. Scale bars, 100 µm.

All results are from day 56 of HI and expressed as mean±SEM, relative to the isogenic control (dotted lines). Where not otherwise indicated, samples are n=4 biological replicates. As experiments were performed at the same time, data of untreated mutants and isogenic controls in (A) to (C) are the same as in Figures 3C, 3D, 4A, and 4D. CBZ, carbamazepine; TMAO, trimethylamine N-oxide; VPA, valproic acid; RM, rapamycin. (*p < 0.05, **p < 0.01, ***p < 0.001 by unpaired two-sided t-test of treated compared to untreated samples; indication of n.s., no significance, is omitted).

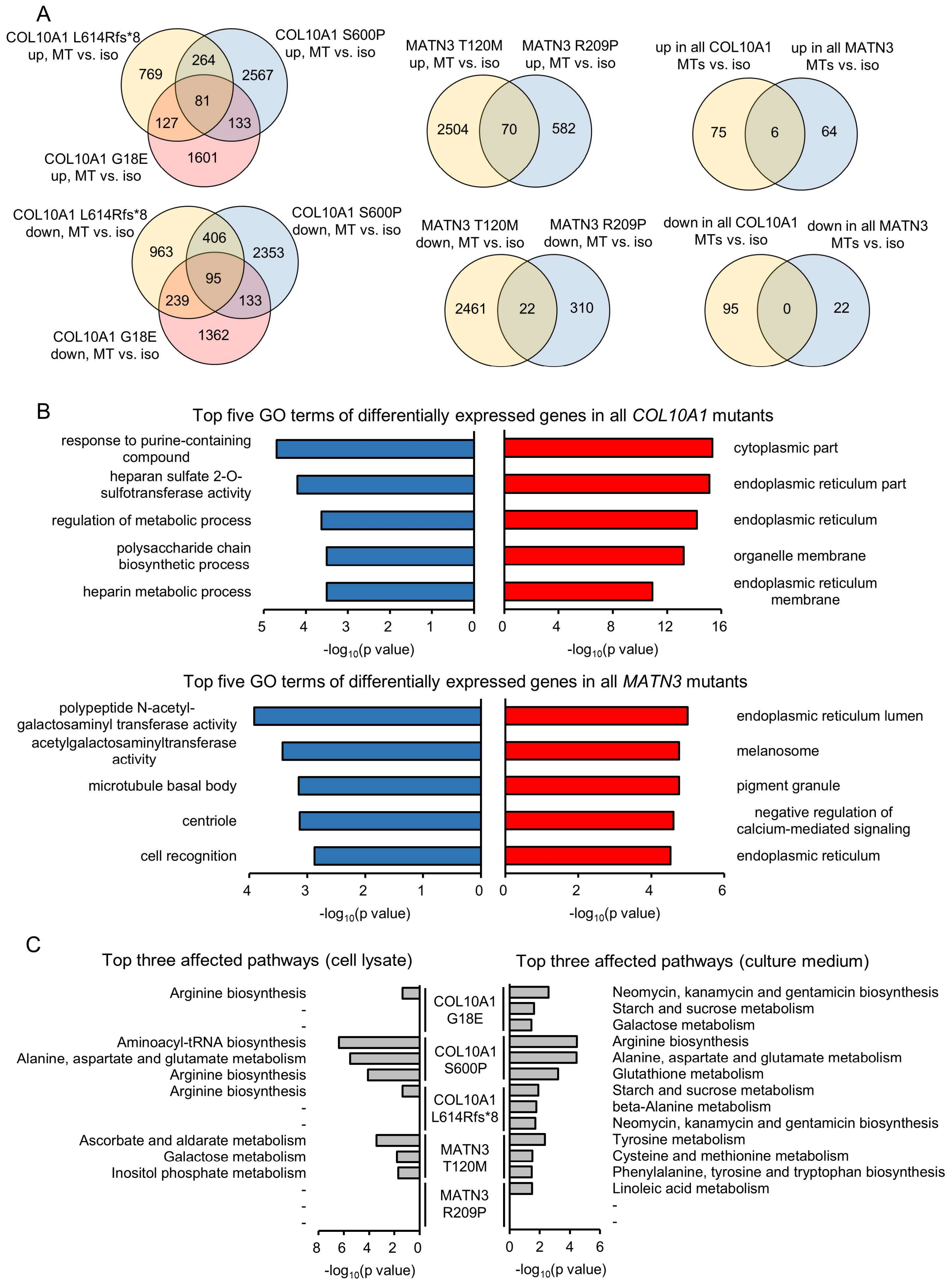

A
COL10A1 L614Rfs*8 up, MT vs. iso
COL10A1 S600P up, MT vs. iso
COL10A1 G18E up, MT vs. iso
769
264
2567
81
127
133
1601
MATN3 T120M up, MT vs. iso
MATN3 R209P up, MT vs. iso
2504
70
582
up in all COL10A1 MTs vs. iso
up in all MATN3 MTs vs. iso
75
6
64
COL10A1 L614Rfs*8 down, MT vs. iso
COL10A1 S600P down, MT vs. iso
COL10A1 G18E down, MT vs. iso
963
406
2353
95
239
133
1362
MATN3 T120M down, MT vs. iso
MATN3 R209P down, MT vs. iso
2461
22
310
down in all COL10A1 MTs vs. iso
down in all MATN3 MTs vs. iso
95
0
22
B
Top five GO terms of differentially expressed genes in all COL10A1 mutants
response to purine-containing compound
heparan sulfate 2-O-sulfotransferase activity
regulation of metabolic process
polysaccharide chain biosynthetic process
heparin metabolic process
cytoplasmic part
endoplasmic reticulum part
endoplasmic reticulum
organelle membrane
endoplasmic reticulum membrane
5 4 3 2 1 0
0 4 8 12 16
-log10(p value)
-log10(p value)
Top five GO terms of differentially expressed genes in all MATN3 mutants
polypeptide N-acetyl-galactosaminyl transferase activity
acetylgalactosaminyltransferase activity
microtubule basal body
centriole
cell recognition
endoplasmic reticulum lumen
melanosome
pigment granule
negative regulation of calcium-mediated signaling
endoplasmic reticulum
4 3 2 1 0
0 2 4 6
-log10(p value)
-log10(p value)
C
Top three affected pathways (cell lysate)
Top three affected pathways (culture medium)
COL10A1 G18E
Arginine biosynthesis
-
-
Neomycin, kanamycin and gentamicin biosynthesis
Starch and sucrose metabolism
Galactose metabolism
COL10A1 S600P
Aminoacyl-tRNA biosynthesis
Alanine, aspartate and glutamate metabolism
Arginine biosynthesis
Arginine biosynthesis
Alanine, aspartate and glutamate metabolism
Glutathione metabolism
COL10A1 L614Rfs*8
Arginine biosynthesis
-
-
Starch and sucrose metabolism
beta-Alanine metabolism
Neomycin, kanamycin and gentamicin biosynthesis
MATN3 T120M
Ascorbate and aldarate metabolism
Galactose metabolism
Inositol phosphate metabolism
Tyrosine metabolism
Cysteine and methionine metabolism
Phenylalanine, tyrosine and tryptophan biosynthesis
MATN3 R209P
-
-
-
Linoleic acid metabolism
-
-
8 6 4 2 0
0 2 4 6
-log10(p value)
-log10(p value)

**Figure S7. Commonly and differently affected pathways in *COL10A1* and *MATN3* mutants. Related to Figure 7**

(A) Venn diagram of commonly upregulated (top) and downregulated (bottom) genes in each mutant (MT) compared to its isogenic control (iso) with $p<0.05$ by moderated t-test. Genes in the central sections of the left and middle Venn diagrams were used for GO analysis in (B).

(B) Top five GO terms of differentially expressed genes in all *COL10A1* (top) or all *MATN3* (bottom) mutants compared to their isogenic controls. Red, upregulated; blue, downregulated in the mutant.

(C) Top three affected pathways of differentially abundant metabolites in each clone calculated from the cell lysate (left) and culture medium (right). Statistical analysis was performed using a two-sided t-test with unequal variances, and metabolites with $p<0.05$ were included in the pathway analysis. The legend shows (-) if fewer than three pathways were detected for that sample.

All results are from n=3 biological replicates from day 56 of HI.

**Supplemental Tables**

**Table S1. Number of biological replicates for mRNA expression in Figures 2B, 4A, 6A and Supplementary Figures S5A, S5E, S6A**

| Sample name | Mutant or isogenic control | Number of biological replicates |
|---|---|---|
| COL10A1 G18E | mutant (#1) | 12 |
| | mutant #2 | 4 |
| | isogenic control (#1) | 8 |
| COL10A1 S600P | mutant (#1) | 12 |
| | mutant #2 | 4 |
| | isogenic control (#1) | 12 |
| COL10A1 L614Rfs*8 | mutant (#1) | 8 |
| | mutant #2 | 4 |
| | isogenic control (#1) | 4 |
| | isogenic control #2 | 4 |
| MATN3 T120M | mutant (#1) | 4 |
| | mutant #2 | 4 |
| | isogenic control (#1) | 4 |
| | isogenic control #2 | 4 |
| MATN3 R209P | mutant (#1) | 8 |
| | mutant #2 | 4 |
| | isogenic control (#1) | 12 |
| All drug-treated groups | - | 4 |

[a]Unless stated otherwise, all mutants and isogenic controls in the main and supplemental figures refer to mutants #1 and isogenic controls #1.

**Table S2. List of differentially expressed genes in the heatmap of Figure 7D**

| Endoplasmic reticulum GO:0005783 | Protein transport GO:0015031 | Proteasome complex GO:0000502 | Lysosome GO:0005764 | Regulation of metabolic process GO:0019222 | Induction of apoptosis GO:0006917 |
|---|---|---|---|---|---|
| HSP90B1 | SLC15A4 | PSMA5 | CTSF | TGFA | MUL1 |
| PDIA4 | GPAM | PSMB3 | NPC2 | IRX6 | LTA |
| HSPA5 | GPAM | PSMC6 | CHID1 | PLA2G10 | LTA |
| DERL3 | GPAM | PSMC6 | PLA2G15 | ZNF253\|ZNF626\|ZNF93 | PPP1R13B |
| WFS1 | MIA3 | VCP | DNASE2 | TADA2A | LTA |
| PDIA6 | EXPH5 | PSMB7 | HYAL2 | PRDM5 | TGFBR1 |
| CRELD2 | GRIK2 | PSMD14 | KCNE2 | PPARG | DLC1 |
| EPHX1 | TGFB1 | PSMA3 | ITM2C | DEPDC6 | XPA |
| LPCAT3 | EIF5AL1 | PSMD12 | PCYOX1 | PRKAA2 | PRKCE |
| MLEC | EIF5AL1 | PSMB4 | GUSB | ZNF774 | TFB1M\|TIAM2 |
| KDELC2 | EIF5AL1 | PAAF1 | TM9SF1 | ZEB2 | IFI16 |
| DPAGT1 | SEC23IP | PSMA1 | NEU4 | EGR3 | SMAD3 |
| TMEM8B | ARF4 | PSMA4 | MANBA | VAV2 | C16orf5 |
| HLA-DPA1 | XPO7 | PSMB11 | ABCA2 | GNG7 | IKBKE\|C1orf147 |
| HM13 | PITPNM1 | PSMB2 | CTBS | YEATS2 | ARHGAP4 |
| MPPE1 | AKAP5 | POMP | IFI30 | ZNF585B | DEDD |
| SEC22B | TMCO6 | PSMD8 | FNBP1 | GMFG | YJEFN3\|NDUFA13 |
| SMPD4 | LMAN1 | PSMD10 | NPC1 | ZNF454 | AIFM3 |
| G6PC3 | SEC22B | PSMC4 | PPT1 | ADNP | TLR2 |
| EIF5AL1 | SRPR | PSMC3 | DRAM2 | CABYR | ITM2B |
| EIF5AL1 | ARFGAP1 | PSMF1 | LMBRD1 | ZBTB10 | LTB4R2\|CIDEB\|C14orf21 |
| EIF5AL1 | KDELR2 | ADRM1 | CTSB | HNRNPF | BRCA1 |
| HLA-DPB1 | COPE | PSMD11 | HLA-DQB1\|HLA-DRB1\|LOC100133583\|LOC100509 | SMURF2 | BCL2L13 |

| | | | | | |
|---|---|---|---|---|---|
| | | | 325 | | |
| HLA-DPB1 | KDELR1 | IDE | C1orf85 | ZNF266 | PHLDA1 |
| MIA3 | SCFD2 | PSMD3 | CTSL1 | SIK1 | FAS |
| DNAJC10 | YIF1A | PSME3 | VAMP4 | TIGD6\|HMGXB3 | BAX |
| ERAP1 | KDELR3 | PSMD1 | GBA | VEZF1 | IKBKG |
| TPD52 | CHMP4C | PSMC2 | NAGA | ACER2 | BNIP3 |
| SERP1 | COPZ1 | | OR10V1\|STX3 | TAF4B | PPARG |
| CLCC1\|GPSM2 | BET1L | | PLBD2 | RASGRF2 | ITGB3BP |
| PREB | PEX6 | | UNC93B1 | NHEJ1\|SLC23A3 | VAV2 |
| ERLEC1 | IPO13 | | SCARB2 | GTF2IRD2B | THBS1 |
| STT3A | ARF3 | | AZU1 | GTF2IRD2\|GTF2IRD2B | AHRR\|PDCD6 |
| SERPINH1 | RAB40B | | UNC93B1\|UNC93B5 | SPRY2 | PLAGL2 |
| RPN1 | BLOC1S3 | | LIPA\|LOC100509637 | ARHGEF3 | RASGRF2 |
| | VPS39 | | CD164 | P2RY2 | ARHGEF3 |
| | COPG | | HLA-DRB3\|HLA-DRB1 | CAMK2A | DAPK1 |
| | NSF | | HLA-DRB4\|HLA-DRB1 | DOCK4 | INHBA |
| | COPB2 | | CD74 | INPP5D | PRUNE2 |
| | RAB43 | | HLA-DMA\|HLA-DMB | TSHZ2 | ERN1 |
| | AP3D1 | | HLA-DMA\|HLA-DMB | TSHZ2 | ERCC6\|PGBD3 |
| | RRBP1 | | HLA-DMA\|HLA-DMB | KAL1 | CUL2 |
| | SEC61A1 | | STS | | CIDECP |
| | KATNB1 | | IDS | | HIPK1 |
| | PEX10\|RER1 | | LGMN | | TRAF3 |

| | | | | | |
|---|---|---|---|---|---|
| | VPS4A | | RAB14 | | SAP30BP |
| | CDC37\|MIR1181 | | IDS | | SOS2 |
| | TOMM40 | | HLA-DPB1 | | ERCC3 |
| | YKT6 | | HLA-DPB1 | | CEBPG |
| | HSP90B1 | | SMPD1 | | DAPK2 |
| | PDIA4 | | SLC17A5 | | YWHAE\|PAFAH1B1 |
| | GOPC\|DCBLD1\|ROS1 | | CD68 | | PTEN |
| | HERC2P2\|HERC2P3\|HERC2P9 | | HLA-DMB | | SENP1\|PFKM |
| | HERC2P3\|HERC2P2\|HERC2P9 | | TMEM55B | | PTENP1 |
| | STEAP3 | | ANKRD27 | | ARHGEF7 |
| | PREB | | PRCP | | FOXO3B |
| | SERP1 | | HYAL1 | | AIFM2 |
| | RAB25 | | GM2A | | TNFRSF10B |
| | RAB26 | | ARSB | | AEN |
| | MVP | | TRIP10 | | RIPK1 |
| | APOC2 | | HLA-DPA1 | | VAV3 |
| | CD24 | | HLA-DPA1 | | ARHGEF12 |
| | VPS37B | | HLA-DPA1 | | SH3RF1 |
| | AKT1 | | CCKAR | | SOS1 |
| | PDIA3P | | LDLR | | ARHGEF2 |
| | PDIA3 | | GPC3 | | STK17A |
| | CALR | | | | TP53I3 |
| | LIN7B | | | | ZMAT3 |
| | SNX21 | | | | TUBB2C |
| | | | | | ABR |
| | | | | | BNIP3 |
| | | | | | YWHAE |

[a]Genes are listed in the order of appearance in Figure 7D.

[b]Genes listed more than once have different Transcript Cluster IDs.

**Table S3. List of differentially abundant metabolites in *COL10A1* mutants in Figure 7E**

| | Cell lysate | | Culture medium | |
|---|---|---|---|---|
| | Compound name | FC | Compound name | FC |
| COL10A1 G18E | Arachidonic acid | MT only | Pimelic acid | 2.10 |
| | Ornithine | 4.78 | Glucose | 0.45 |
| | 3-Phosphoglyceric acid | 0.61 | | |
| | Dihydroxyacetone phosphate | 0.28 | | |
| | 3-Hydroxyisobutyric acid | iso only | | |
| COL10A1 S600P | Succinylacetone-ox-origin fragment | MT only | Cystine | 5.38 |
| | 5-Dehydroquinic acid | MT only | Succinic acid | 1.45 |
| | Dihydroxyacetone phosphate | 2.97 | Malic acid | 1.25 |
| | Hexanoylglycine | 2.81 | 2-Hydroxyglutaric acid | 1.21 |
| | 4-Aminobutyric acid | 2.60 | 3-Hydroxyglutaric acid | 1.21 |
| | Ribose | 1.80 | Guanine | 1.08 |
| | 3-Dehydroshikimic acid | 1.70 | Glucono-1,5-lactone | 0.98 |
| | 2-Hydroxyadipic acid | 1.32 | Fructose | 0.85 |
| | 6-Phosphogluconic acid | 0.90 | Sorbose | 0.85 |
| | Inositol phosphate | 0.90 | Glyceric acid | 0.84 |
| | N-Acetylaspartic acid | 0.77 | Galacturonic acid | 0.65 |
| | N-Acetylaspartic acid | 0.75 | Lyxose | 0.64 |
| | Methoprene acid | 0.74 | Arabinose | 0.64 |
| | Inositol | 0.74 | 5-Oxoproline | 0.61 |
| | N-Acetylmannosamine | 0.73 | 2-Ketoglutaric acid | 0.61 |
| | Rhamnose | 0.71 | Cysteine | 0.58 |
| | Leucine | 0.70 | Ribose | 0.57 |
| | Oxalic acid | 0.69 | Propionylglycine | 0.56 |
| | 2-Hydroxybutyric acid | 0.69 | Aspartic acid | 0.56 |
| | Tyrosine | 0.69 | Glutamic acid | 0.56 |
| | Mannitol | 0.69 | Linoleic acid | 0.51 |
| | Ethylmalonic acid | 0.68 | Glucose 6-phosphate | 0.50 |
| | Glucose | 0.68 | Hypoxanthine | 0.47 |
| | Glucosamine | 0.68 | Ornithine | 0.45 |
| | Mannose | 0.67 | Glycyl-Glycine | 0.45 |
| | Galactose | 0.67 | Decanoic acid | 0.35 |
| | Glucose | 0.67 | 2-Ketoisocaproic acid | 0.25 |
| | Lysine | 0.66 | Pyruvic acid | 0.24 |
| | Glycine | 0.65 | 3-Methyl-2-oxovaleric acid | 0.15 |
| | Isobutyrylglycine | 0.65 | Octanoic acid | iso only |
| | O-Acetylserine | 0.65 | | |
| | 3-Aminopropanoic acid | 0.65 | | |
| | 1,6-Anhydroglucose | 0.65 | | |
| | Threonine | 0.64 | | |

| | | |
|---|---|---|
| | Ureidosuccinic acid | 0.63 |
| | Octanoic acid | 0.61 |
| | Ribulose | 0.61 |
| | Xylulose | 0.61 |
| | Glucono-1,4-lactone | 0.60 |
| | Arginine | 0.58 |
| | Tryptophan | 0.58 |
| | Adenosine monophosphate | 0.58 |
| | 2-Propyl-3-hydroxy-pentanoic acid | 0.56 |
| | 2-Propyl-3-hydroxy-pentanoic acid | 0.56 |
| | 5-Oxoproline | 0.55 |
| | Asparagine | 0.55 |
| | Xanthosine monophosphate | 0.54 |
| | Cadaverine | 0.53 |
| | Hypotaurine | 0.52 |
| | Aspartic acid | 0.50 |
| | Fumaric acid | 0.46 |
| | Propionylglycine | 0.46 |
| | 2-Hydroxyglutaric acid | 0.45 |
| | 3-Hydroxyglutaric acid | 0.45 |
| | Glutamic acid | 0.45 |
| | Serine | 0.44 |
| | Galacturonic acid | 0.44 |
| | Galactose | 0.41 |
| | Xylose | 0.39 |
| | Galactose | 0.38 |
| | Tryptamine | 0.38 |
| | Glucuronic acid | 0.35 |
| | Galactosamine | 0.34 |
| | Tagatose | 0.29 |
| | Dihydrouracil | 0.27 |
| | Succinylacetone | 0.27 |
| | O-Phosphoethanolamine | 0.27 |
| | 2-Ketoisocaproic acid | 0.26 |
| | Benzoic acid | 0.26 |
| | 2-Ketoglutaric acid | 0.26 |
| | 2-Ketoglutaric acid | 0.25 |
| | 3-Methyl-2-oxovaleric acid | 0.23 |
| | 4-Hydroxybutyric acid | 0.23 |
| | Dihydroxyacetone | 0.22 |
| | 2-Ketoisocaproic acid | 0.21 |
| | Dopamine | 0.21 |

| | | | | |
|---|---|---|---|---|
| | Spermidine | 0.20 | | |
| | Acetylglycine | 0.16 | | |
| | 3-Methyl-2-oxovaleric acid | 0.15 | | |
| | N-Acetyltyrosine | 0.10 | | |
| | Biotin | 0.05 | | |
| | Glucose | iso only | | |
| COL10A1 L614Rfs*8 | Octenedioic acid | MT only | 3-Methylglutaconic acid | MT only |
| | Ornithine | 3.92 | 2-Aminoadipic acid | MT only |
| | Palmitoleic acid | 2.48 | Fructose | MT only |
| | Pimelic acid | 1.87 | 3-Hydroxydodecanedioic acid | MT only |
| | Saccharopine | 0.36 | Glucose | 12.16 |
| | Ribose | 0.12 | Galactosamine | 7.21 |
| | | | Galacturonic acid | 6.42 |
| | | | Tryptamine | 6.30 |
| | | | Dopamine | 2.71 |
| | | | Boric acid | 2.27 |
| | | | Dihydrouracil | 2.15 |
| | | | Spermine | 2.11 |
| | | | Cystamine-nTMS | 1.77 |
| | | | 4-Hydroxybutyric acid | 1.54 |
| | | | N-Acetylserine | 1.45 |
| | | | Uridine | 1.38 |
| | | | Glycolic acid | 1.34 |
| | | | Proline | 1.21 |
| | | | Lactitol | 1.17 |
| | | | Maltose | 1.17 |
| | | | Sorbose | 0.70 |
| | | | Ascorbic acid | 0.57 |
| | | | 2-Hydroxyadipic acid | 0.27 |

[a]FC indicates the fold change of mutant vs. isogenic control.

[b]"MT only" or "iso only" indicate the presence of a metabolite only in the mutant (MT) or isogenic control (iso).

[c]Metabolites that are listed more than once have different amounts of TMS.

**Table S4. List of differentially abundant metabolites in *MATN3* mutants in Figure 7E**

| | Cell lysate | | Culture medium | |
|---|---|---|---|---|
| | Compound name | FC | Compound name | FC |
| MATN3 T120M | Saccharopine | 2.41 | 2-Hydroxyisovaleric acid | 2.95 |
| | 3-Dehydroshikimic acid | 2.35 | 2-Hydroxyglutaric acid | 1.99 |
| | Coniferyl aldehyde | 2.22 | 3-Hydroxyglutaric acid | 1.99 |
| | Galacturonic acid | 2.20 | Pyridoxamine | 1.94 |
| | Sorbitol | 2.19 | Octopamine | 1.91 |
| | Galactitol | 2.19 | Cystine | 1.66 |
| | meso-Erythritol | 2.10 | Guanine | 1.54 |
| | Ascorbic acid | 2.07 | meso-Erythritol | 1.50 |
| | Dihydroxyacetone phosphate | 1.93 | Uridine | 1.47 |
| | Glucaric acid | 1.86 | 4-Hydroxyphenylpyruvic acid | 1.45 |
| | Proline | 1.74 | Pyruvic acid | 1.43 |
| | Glucuronic acid lactone | 1.70 | Homoserine | 1.41 |
| | Pimelic acid | 1.64 | 6-Phosphogluconic acid | 1.41 |
| | 3-Hydroxyisobutyric acid | 1.62 | Inositol phosphate | 1.40 |
| | Methoprene acid | 1.61 | 2-Hydroxyisobutyric acid | 1.36 |
| | N-Acetylmannosamine | 1.58 | Dopamine | 0.65 |
| | Pantothenic acid | 1.56 | Decanoic acid | 0.46 |
| | 6-Phosphogluconic acid | 1.56 | 2-Aminopimelic acid | 0.34 |
| | Inositol phosphate | 1.56 | | |
| | 4-Hydroxyphenyllactic acid | 1.52 | | |
| | Uridine monophosphate | 1.51 | | |
| | Niacinamide | 1.50 | | |
| | Guanine | 1.47 | | |
| | Uridine | 1.46 | | |
| | Inositol | 1.43 | | |
| | Ribonolactone | 1.36 | | |
| | Glycine | 1.35 | | |
| | Cholesterol | 1.27 | | |
| | Lactic acid | 1.22 | | |
| | Tagatose | 0.81 | | |
| | Succinylacetone | 0.59 | | |
| | Psicose | 0.59 | | |
| | Fructose | 0.59 | | |
| | Dihydrouracil | 0.56 | | |
| | Citric acid | 0.42 | | |
| | Arachidonic acid | 0.40 | | |
| MATN3 R209P | Hexanoylglycine | 5.06 | 3,6-Epoxydodecanedioic acid | 2.45 |
| | Boric acid | 1.59 | Inosine | 2.10 |
| | Pyridoxal | 0.73 | 3-Sulfinoalanine | 2.04 |
| | O-Phosphoethanolamine | 0.49 | Glutaconic acid | 1.76 |

| | | | | |
|---|---|---|---|---|
| | 4-Hydroxybutyric acid | 0.41 | Ribose | 1.75 |
| | Acetylglycine | 0.38 | Fucose | 1.75 |
| | Dopamine | 0.33 | Palmitoleic acid | 1.69 |
| | Benzoic acid | 0.32 | Linoleic acid | 1.29 |
| | N-Acetyltyrosine | 0.21 | Pimelic acid | 1.25 |
| | Fucose | 0.20 | Succinic acid | 0.67 |
| | Mannitol | 0.04 | Lyxose | 0.61 |
| | Succinic acid | iso only | Glutaric acid | 0.53 |
| | | | Pyridoxal | iso only |
| | | | 3-Methylglutaconic acid(Z) | iso only |

[a]FC indicates the fold change of mutant vs. isogenic control.

[b]"MT only" or "iso only" indicate the presence of a metabolite only in the mutant (MT) or isogenic control (iso).

[c]Metabolites that are listed more than once have different amounts of TMS.

**Table S5. Guide RNAs and repair templates used for gene editing. Related to Experimental Procedures**

| Gene editing | Guide RNA (protospacer) | Single-stranded oligonucleotide repair template (100 nt) | Restriction enzyme |
|---|---|---|---|
| Creation of COL10A1 S600P | CGTGCATGTGAAAGGGACTC | CCCAAGGACTGGAATCTTTACTTGTCAGATACCAGGAATATACTATTTTCCATATCACGTACATGTGAAAGGGACTCATGTTTGGGTAGGCCTGTATAAG | PciI (ACATGT) |
| Creation of MATN3 R209P | AGGTGAATGAAGTGGCGGCT | ATCATTGTTACAGATGGGAGGCCCCAGGACCAGGTGAATGAAGTGGCCGCTCCGGCCCAAGCATCTGGTATTGAGCTCTATGCTGTGGGCGTGGACCGGG | BsrBI (CCGCTC) |
| Rescue of COL10A1 G18E | TGAACTTGGTTCATGAAGTG | AGAATATGCTGCCACAAATACCCTTTTTGCTGCTAGTATCATTGAACCTGGTCCATGGAGTGTTTTACGCTGAACGATACCAAATGCCCACAGGCATAAA | NcoI (CCATGG) |
| Rescue of COL10A1 L614Rfs*8 | TTGGGTAGGCCGTATAAGAA | TTCATACCACGTGCATGTGAAAGGGACTCATGTTTGGGTAGGCCTTTATAAGAATGGCACCCCTGTAATGTACACCTATGATGAATACACCAAAGGCTAC | PsiI (TTATAA) |
| Rescue of MATN3 T120M | ACATGCGGGTGGCAGTGGTG | AACTTTTGTCTCCCGGATAATCGACACTCTGGACATTGGGCCAGCTGACACGCGGGTGGCAGTGGTGAACTATGCTAGCACTGTGAAGATCGAGTTCCAA | PvuII (CAGCTG) |

[a]Underlined nucleotides in the repair template indicate the recognition sites of the restriction enzymes.

**Table S6. Primers for qPCR and sequencing. Related to Experimental Procedures**

| Used in | Amplified gene/domain | Direction | Sequence |
|---|---|---|---|
| sequencing | COL10A1 NC1 domain | forward | CAGCAGGAGCAAAGGGAATG |
| | | forward | GCTGGCATAGCAACTAAGG |
| | | forward | CCACCAGGTCAAGCAGTCATG |
| | | reverse | TTGAATGGGAGGCACAAGG |
| | | reverse | GGGAAGGTTTGTTGGTCTG |
| | | reverse | TGTACTCACATTGGAGCCACTAG |
| | COL10A1 NC2 domain | forward | TCAGTCTGTGGATGATAGTC |
| | | forward | CACTTGATTACCTGAGTATAGC |
| | | reverse | AGTTTCATTTGCCTGCTTG |
| | | reverse | CAGAAGTTGGAAAGTAACACC |
| | MATN3 vWFa domain | forward | GCAGAGGGATGAGGTTCTAAG |
| | | forward | GCAGGCAGTGGTGATGTTGG |
| | | reverse | GTTGGAAGCAAAGACTGACC |
| | | reverse | GGCTTCGTCCATTGCTGTCTG |
| qPCR | ACTB | forward | CACCATTGGCAATGAGCGGTTC |
| | | reverse | AGGTCTTTGCGGATGTCCACGT |
| | COL10A1 | forward | CCCAGCACGCAGAATCCATC |
| | | reverse | AGTGGGCCTTTTATGCCTGT |
| | COL2A1 | forward | CGAGGCAACGATGGTCAGCC |
| | | reverse | TGGGGCCTTGTTCACCTTTGA |
| | MATN3 | forward | GTTCAGTCCGTGACAAGTGTG |
| | | reverse | AGTCTTCGTGCTTCCTCAGTG |
| | SOX9 | forward | GACTTCCGCGACGTGGAC |
| | | reverse | GTTGGGCGGCAGGTACTG |
| | MMP13 | forward | CATGAGTTCGGCCACTCCTT |
| | | reverse | CCTGGACCATAGAGAGACTGGA |
| | IHH | forward | CGGTGGACATCACCACATCA |
| | | reverse | CGTGGGCCTTTGACTCGTAA |
| | ALPL | forward | GGAAGACACTCTGACCGTGG |
| | | reverse | GGGGGCCAGACCAAAGATAG |
| | SP7 | forward | ATCCAGCCCCCTTTACAAGC |
| | | reverse | TAGCATAGCCTGAGGTGGGT |
| | ACAN | forward | TCGAGGACAGCGAGGCC |
| | | reverse | TCGAGGGTGTAGCGTGTAGAGA |
| | RUNX2 | forward | TTACTTACACCCCGCCAGTC |
| | | reverse | TATGGAGTGCTGCTGGTCTG |

| | | | |
|---|---|---|---|
| | COL9A2 | forward | GCGCTATCGGTGCCACTGGG |
| | | reverse | GGGGGCCCGTTGCTCCTTTC |
| | HSPA5 | forward | GTTTGCTGAGGAAGACAAAAAGCTC |
| | | reverse | CACTTCCATAGAGTTTGCTGATAATTG |
| | CRELD2 | forward | CCAAGTACGAGTCCAGCGAG |
| | | reverse | TGCTCCTCCTGCGCCTCTAG |
| | HSP90B1 | forward | ATGGAGCAGCAAGACTGAAAC |
| | | reverse | TCTTCTTCTTCTTCCTCTACTGC |

**Table S7. Key reagents used in this study. Related to Experimental Procedures**

| Sclerotome induction reagents | Source | Identifier |
|---|---|---|
| AK02N | Ajinomoto | AJ100 |
| iMatrix-511 silk | Nippi | 892021 |
| Matrigel | BD | 354230 |
| PenStrep | Gibco | 15140-163 |
| bFGF | Wako | 068-04544 |
| CHIR99021 | Axon | Axon1386 |
| Activin A | R&D | 338-AC01-M |
| SB431542 | Selleck Chemicals | S1067 |
| LDN193189 | Stemgent | 04-0074 |
| PD173074 | Tocris | 3044 |
| XAV939 | Tocris | 3748/10 |
| SAG | Calbio | 566660 |
| Y-27632 | Wako | 034-24024 |
| Ham's F12 | Gibco | 11765-054 |
| BSA | Sigma | A8806-5G |
| IMDM | Sigma | I3390 |
| CD Lipid | Life Tech | 11905-031 |
| Apo-transferrin | Sigma | T1147-500MG |
| Monothioglycerol | Sigma | M6145-25ML |
| Insulin | Wako | 097-06474 |
| Hypertrophic induction reagents | Source | Identifier |
| Dexamethasone | Wako | 047-18863 |
| PDGF-BB | R&D | 520-BB |
| TGFβ3 | R&D | 243-B3 |
| BMP4 | R&D | 314-BP |
| T3 | Sigma | T-074-1ML |
| β-glycerophosphate | Sigma | G6501-100G |
| ITS premix | Corning | 354352 |
| L-ascorbic acid 2-phosphate | Sigma | A8960 |
| Proline | Sigma | P-5607 |
| Glucose | Sigma | G8769 |

| | | |
|---|---|---|
| Sodium pyruvate | Sigma | S8636 |
| Glutamax | Life Tech | 35050-061 |
| DMEM/F12 | Gibco | 11320-033 |
| Autophagy inducers and chemical chaperones | Source | Identifier |
| Carbamazepine | Sigma | C4024-1G |
| Trimethylamine N-oxide | Sigma | 317594-5G |
| Valproic acid | Sigma | P4543-10G |
| Rapamycin | MedChemExpress | HY10219 |
| Antibodies (Simple Western) | Source | Identifier |
| ACTB | CST | 4970S |
| HSPA5 | CST | 3177T |
| PDI | CST | 3501T |
| Antibodies (immunofluorescence) | Source | Identifier |
| COL2A1 | Thermo | MS235P0 |
| COL10A1 | Thermo | 14-9771-82 |
| MATN3 | abcam | ab106388 |
| PDI | Thermo | MA3-019-A647 |
| Antibodies (FACS analysis) | Source | Identifier |
| DLL1 | R&D | FAB1818A |
| APC-conjugated mouse IgG2B | R&D | IC0041A |
| Antibodies (iPSC validation) | Source | Identifier |
| SSEA4 | Chemicon | MAB4304 |
| TRA1-60 | Chemicon | MAB4360 |
| TRA1-81 | Chemicon | MAB4381 |

**Supplemental Experimental Procedures**

**Establishment of isogenic iPSC lines using CRISPR/Cas9**

In order to create gene-corrected rescues, the 20 nt protospacer sequence for the guide RNA targeting the mutated site was inserted into the vector pSpCas9(BB)-2A-Puro (PX459) V2.0 (Addgene). Cloning was performed as previously described (Ran et al., 2013) using DH5α cells (Toyobo). The vector was isolated using the Midi Prep Kit (Macherey-Nagel) and electroporated into iPSCs ($1 \times 10^6$ cells) using the NEPA 21 (Nepa Gene), together with a repair template consisting of 100 nt including the gene correction, a recognition site for restriction enzymes to confirm the insertion, and silent mutations for PAM or guide RNA blocking. Cells were cultured in six well plates with 10 μM Y-27632 overnight, after which they were treated with 0.5-0.7 μg/ml puromycin for 24 hours to select for cells incorporating the vector. One week after sparsely re-seeding the cells, 48-96 single colonies were randomly picked and expanded. Genomic DNA was extracted using the DNeasy Blood and Tissue Kit (QIAGEN) with proteinase K treatment, upon which the relevant regions were amplified with PCR using KOD Plus Neo (Toyobo). Amplicons were digested by the restriction enzymes to confirm insertion of the repair template. Samples for which insertion was confirmed were then purified using the FastGene Gel/PCR Extraction kit (Nippon Genetics) and sequenced using the BigDye Terminator v3.1 Cycle Sequencing Kit (Applied Biosystems) with the 3500xL Genetic Analyzer (Applied Biosystems).

Mutants were established from the wild type iPSC line 414C2 similarly to the gene-corrected rescues. Here, the guide RNA was designed to target a SNP near the intended site of mutagenesis. The 100 nt repair template included the new mutation, a recognition site for restriction enzymes, and silent mutations for PAM or guide RNA blocking. All primers, guide RNAs, repair templates, and restriction enzymes used in this gene editing are listed in Tables S5 and S6.

**iPSC culture and validation**

All patient-derived iPSC lines were established on SNL feeder cells in primate embryonic stem cell medium (ReproCELL) supplemented with 4 ng/ml bFGF (Wako), 50 U penicillin and 50 μg/ml streptomycin (Gibco). After several passages as whole colonies on feeder cells, iPSCs were moved to feeder-free culture on dishes coated with laminin (Nippi) in StemFit AK02N (Ajinomoto) and passaged as single cells once a week at a density of $1.1 \times 10^3$ to $3.2 \times 10^3$ cells/$cm^2$.

Newly established iPSC clones were validated for pluripotency by immunostaining either before switching to feeder-free culture (MATN3 T120M and COL10A1 L614Rfs*8) or after the switch (COL10A1 G18E). After fixation in 4% paraformaldehyde, cells were PBS washed and incubated for 1 hour at 4°C in a blocking solution of PBS with 0.2% TritronX-100 (Sigma) and 5% Blocking One Histo (Nacalai Tesque). Pluripotency markers SSEA4, TRA1-60, and TRA1-81 (all Chemicon) were diluted in the blocking solution at 1:400, 1:1000, and

1:1000, respectively, and used to stain the cells overnight at 4°C. On the next day, cells were further incubated for 1 hour at 4°C in blocking solution containing Alexa 488-conjugated goat anti-mouse IgG antibody (Thermo) at 1:500. The nucleus was counterstained using DAPI.

All iPSC clones were further validated for their ability to differentiate into all three germ layers through teratoma formation. $1\times10^6$ cells were injected into each testis of at least three NOD-SCID mice, which were sacrificed after 8 to 12 weeks to excise the tissue. After fixation in 4% paraformaldehyde, the tissue was embedded in paraffin, sectioned, and stained with H&E.

**Sclerotome induction (SI) and hypertrophic induction (HI)**

For SI, CDMi (chemically defined medium with insulin) was used as base medium (Wataya et al., 2008). This medium consists of a 1:1 mixture of Iscove's modified Dulbecco's medium (IMDM) (Sigma) and Ham's F12 (Gibco), 5 mg/ml BSA (Sigma), 1x CD Lipid (Life Tech), 15 µg/ml apo-transferrin (Sigma), 450 µM monothioglycerol (Sigma), 50 U penicillin, 50 µg/ml streptomycin, and 7 µg/ml insulin (Wako). On day 0 of induction, primitive streak cells were induced with 20 ng/ml bFGF, 10 µM CHIR99021 (Axon), and 50 ng/ml Activin A (R&D) for 24 hours. Then, presomitic mesoderm cells were induced with 10 µM SB431542 (Selleck Chemicals), 3 µM CHIR99021, 250 nM LDN193189 (Stemgent), and 20 ng/ml bFGF for 24 hours. Following this, somatic mesoderm cells were induced with 100 nM PD173074 (Tocris) and 1 µM XAV939 (Tocris) for 24 hours. Finally, sclerotome cells were induced with 100 nM SAG (Calbio) and 600 nM LDN193189 for 72 hours.

For HI, HI base medium was used. This medium consists of 1% (v/v) ITS premix (Corning), 0.17 mM L-ascorbic acid 2-phosphate (Sigma), 0.35 mM proline (Sigma), 0.15% (v/v) glucose (Sigma), 1 mM sodium pyruvate (Sigma), 2 mM Glutamax (Life Tech), 100 U penicillin, and 100 µg/ml streptomycin in DMEM/F12 (Gibco). The medium was changed every 2-3 days.

**Quantitative PCR (qPCR) analysis**

For RNA extraction, cartilage pellets were first powderized using the Multi-beads shocker (Yasui Kikai). Total RNA was extracted using the RNeasy Micro or Mini Kit (QIAGEN) with DNase treatment and reverse transcribed using ReverTra Ace (Toyobo) in a total volume of 20 µL with up to 300 ng of RNA. cDNA was diluted 1:10 and qPCR was performed with 1 µl cDNA in duplicate reactions using the Thunderbird SYBR qPCR Mix (Toyobo) and QuantStudio 12K Flex Real-Time PCR System (Thermo). Primers are listed in Table S6. All results were normalized by the β-actin expression in each sample. For statistical analysis, unpaired two-sided t-tests were performed for comparisons between two groups. For comparisons between more than two groups, ANOVA with post-hoc Tukey HSD was used.

**Nonsense-mediated decay (NMD) detection**

Genomic DNA was extracted from iPSCs and RNA was extracted on day 56 of HI of the COL10A1 L614Rfs*8 mutant and isogenic control. For the cycloheximide (CHX)-treated group, cartilage pellets were treated with 100 μg/ml CHX (Sigma) for 6 hours before RNA extraction. RNA was reverse transcribed with a negative control (no reverse transcriptase), and the cDNA was amplified for the NC1 domain with PCR. The amplicons were digested by the restriction enzyme StuI (NEB), which only recognizes the wild type sequence, and BceAI (NEB), which only recognizes the mutant sequence.

**Protein expression analysis**

On day 56 of HI, cartilage pellets were collected, PBS washed, and stored at -80°C. After adding 100 μl SDS sample buffer with 1% (v/v) protease inhibitor cocktail (Nacalai Tesque), the pellets were homogenized using homogenizer pestles (Kenis) and sonicated until complete dissolution. The samples were centrifuged and the supernatant was collected. Protein concentration was measured using the Pierce BCA Protein Assay Kit (Thermo) and protein expression was detected with Simple Western (Wes) using the 12-230 kDa Separation Module (ProteinSimple) according to the manufacturer's instructions. For each lane, 5 μg of protein was used at a concentration of 1 μg/μl. All results were normalized by the β-actin values of each sample. For statistical analysis, unpaired two-sided t-tests were performed.

**Microarray analysis**

After RNA extraction, RNA quality was confirmed using the RNA 6000 Nano Kit (Agilent Technologies). The RNA was reverse transcribed into cDNA and amplified with the Ambion WT Expression Kit, fragmented and labeled with the GeneChip WT Terminal Labeling and Controls kit, and hybridized to Human Gene 1.0ST Arrays using the GeneChip Hybridization Wash and Stain Kit according to the manufacturer's protocol (Affymetrix). The arrays were scanned and the raw data were imported into GeneSpring GX 14.9 for analysis. The data were normalized using the RMA-16 algorithm and the baseline was adjusted to the median of all samples. A moderated t-test was performed on all samples (n=3), and all genes satisfying the $p<0.05$ cut-off were deemed differentially expressed. Heatmaps were generated in GeneSpring by performing GO analysis of differentially expressed genes in one or more mutants and displaying each mutant's average expression of the genes related to the GO terms. Volcano plots were made in R (version 3.6.3) based on the normalized expression data.

**Flow cytometry analysis**

To determine the induction efficiency of presomitic mesoderm, cells were analyzed by flow cytometry on day 2 of SI. Cells were detached and washed with FACS buffer containing 0.1% (w/v) BSA in PBS. Approximately $5\times10^5$ cells were stained with either APC-conjugated DLL1 antibody (R&D) or APC-conjugated mouse IgG2B (R&D) as isotype at a 1:200 dilution in FACS buffer for 30 minutes at 4°C. Following another wash with FACS buffer, the cells were stained with 1 μg/ml DAPI (R&D) to label dead cells. Analysis was performed using the FACS Aria II and the FACSDiva software (BD). Graphs were created using the FlowJo software. For statistical analysis, unpaired two-sided t-tests were performed.

**Histological analysis**

Paraffin-embedded tissue sections were deparaffinized and stained with H&E, von Kossa, and Safranin O. TUNEL staining was performed using the ApopTag Peroxidase *In Situ* Apoptosis Detection kit (Millipore) with proteinase K treatment according to the manufacturer's instructions. For immunostaining, antigen retrieval was performed by 40 minutes incubation at 37°C in 10 mg/ml hyaluronidase (Sigma) and 15 minutes incubation at 80°C in 1 mM EDTA (Wako). After blocking in 10% FBS (Nichirei) for 1 hour, the tissue was stained at 4°C overnight using antibodies against COL2A1 (Thermo) at 1:1000, COL10A1 (Thermo) at 1:300, and MATN3 (abcam) at 1:500 in Can Get Signal Immunostain Solution B (Toyobo). On the next day, the samples were incubated at room temperature for 1 hour in the same solution with Alexa 488-conjugated goat anti-mouse IgG antibody (Thermo) at 1:1000, Alexa 555-conjugated goat anti-rabbit IgG antibody (Thermo) at 1:1000, Alexa 647-conjugated donkey anti-mouse IgG antibody (Thermo) at 1:1000, and Alexa 647-conjugated PDI antibody (Thermo) at 1:500. DAPI was used as a nuclear counterstain at 10 μg/ml and the samples were observed with the BZ-X810 (Keyence).

**Image analysis and quantification**

All samples were quantified using the BZ-X800 Analyzer software and statistical analysis was performed using unpaired two-sided t-tests. Pellet size was quantified using images taken of live pellets from three technical replicates each of four biological replicates on day 56 of HI with the BZ-X810 (Keyence). The size was calculated by taking the average of the major axis of each pellet.

For quantification of the COL2A1 and TUNEL stainings, four images each were taken of four biologically independent samples at x20 magnification, with the images being taken at the top, bottom, left, and right sides of the pellet, at a depth of approximately 500-1000 μm from the surface to avoid both the necrotic center and the disintegrating periphery. Dead cells of each sample were quantified by counting the number of TUNEL-positive cells per field of view (FOV). Cell size was quantified by selecting the inverse of COL2A1 staining and measuring the average area of the selection.

For the quantification of COL10A1, MATN3, and PDI stainings, three images each were taken of four biologically independent samples at x40 magnification randomly throughout the pellet at a depth of approximately 500-1000 μm from the surface to avoid both the necrotic center and the disintegrating periphery. Intracellular retention of COL10A1 and MATN3 was calculated by dividing the total green fluorescence intensity of the area co-staining with PDI (red) by the total green fluorescence intensity of the whole image. The ER size was quantified from the average area of PDI fluorescence.

**Transmission electron microscopy (TEM)**

On day 56 of HI, pellets were PBS washed and fixed in 2.5% glutaraldehyde for 8 hours at 4°C. After PBS washing and post-fixation in 2% osmium tetroxide for 2 hours, the pellets were dehydrated and embedded in Quetol 812 resin. 1 μm sections were taken and stained with toluidine blue. Ultra-thin sections (120 nm) were taken and stained on grids with uranyl acetate and lead citrate. Images were taken using the H-7500 transmission electron microscope (Hitachi) with the NanoSprint500 (AMT).

**Gas chromatography coupled to mass spectrometry (GC-MS)**

On day 56 of HI, pellets were preserved in liquid nitrogen and the culture medium was preserved at -80°C. Pellets were powderized using the Multi-beads shocker and dissolved in 80% methanol. After centrifugation, the supernatant of all samples was collected and lyophilized. Following trimethylsilyl (TMS) derivatization, the metabolites were separated on a DB-5 column (30 m × 0.25 mm id, film thickness 1.0 mm) (Agilent Technologies) and subjected to mass spectrometry. All data were normalized by the internal standard 2-isopropylmalic acid and the DNA per pellet of each sample. The DNA was measured using the Quant-it Pico Green dsDNA Assay Kit (Thermo Fisher) after dissolving pellets at 60°C overnight in buffered papain from papaya latex (Sigma). Statistical analysis to determine differentially abundant metabolites was performed by a two-sided t-test with unequal variances and pathway analysis was performed with MetaboAnalyst.
